# Induced copy-back RNA synthesis as a novel therapeutic mechanism against RNA viruses

**DOI:** 10.1101/2020.02.12.946558

**Authors:** Richard Janissen, Andrew Woodman, Kuo-Ming Lee, Ibrahim Moustafa, Fiona Fitzgerald, Peng-Nien Huang, Louis Kuijpers, Angela L. Perkins, Daniel A. Harki, Jamie J. Arnold, Belen Solano, Shin-Ru Shih, Craig E. Cameron, Nynke H. Dekker

**Author notes:** These authors contributed equally to this work (Richard Janissen, Andrew Woodman). Corresponding author; Email: N. H. D., C. E. C.

## Abstract

The viral RNA-dependent RNA polymerase (RdRp) is a well-established target for development of broad-spectrum antiviral therapeutics. Incorporation of ribonucleotide analogues by the RdRp will either cause termination of RNA synthesis or mutagenesis of the RNA product. We demonstrated recently that incorporation of a pyrazine-carboxamide ribonucleotide into nascent RNA leads to pausing and backtracking of the elongating RdRp. Here, we provide evidence for the single-stranded RNA product of backtracking serving as an intermediate in RdRp-catalyzed, template-switching reactions. This intermediate is used for both intramolecular template-switching (copy-back RNA synthesis) and intermolecular template-switching (homologous RNA recombination). The use of a magnetic-tweezers platform to monitor RdRp elongation dynamics permitted direct observation of copy-back synthesis and illuminated properties of the RdRp that promote copy-back synthesis, including stability of the RdRp-nascent-RNA complex and the dimensions of the RdRp nucleic-acid-binding channel. In cells, recombination was stimulated by the presence of a pyrazine-carboxamide ribonucleotide. The effect of the drug on recombination was diminished for a recombination-defective virus, but this virus was not resistant to the drug. The discovery that a ribonucleotide analogue can induce copy-back RNA synthesis suggests that this third mechanistic class of compounds may function by promoting formation of defective viral genomes. This study identifies RdRp-catalyzed intra- and intermolecular template switching as a viable new mechanistic target with potentially broad-spectrum appeal.

---

It is still not possible to predict the emergence of viruses capable of founding an epidemic, a fact that has been reconfirmed many times over the past few decades. Since the outbreak of West Nile virus in 1999, one RNA virus after another has emerged, causing significant morbidity and mortality on a global scale^1^. The world is under constant threat of an influenza pandemic^1^. An outbreak of enterovirus D-68 (EV-D68), associated with acute flaccid myelitis in young children, is anticipated in 2020^2^. And with recurrent outbreaks of enterovirus A71 in the Asia-Pacific region come hand-foot-and-mouth disease and severe acute flaccid paralysis^3^. This state of affairs demands the availability of broad-spectrum antiviral therapeutics to address the next unanticipated and/or unknown viral pathogen.

Viral polymerases have emerged as tractable and highly efficacious antiviral targets. The retroviral reverse transcriptase (RT) is an essential component of the therapeutic regimen used to treat human immunodeficiency virus (HIV) infection. Past and present protocols to treat hepatitis C virus infection have included compounds targeting the viral RNA-dependent RNA polymerase (RdRp). To date, clinically approved antiviral nucleotides have functioned either by terminating nucleic acid synthesis (chain terminators) or by increasing mutational load on the viral genome (lethal mutagens)^4,5^. Resistance to antiviral nucleotides usually comes with a fitness cost, thus increasing the long-term utility of these classes of antiviral agents ^6^. The major obstacle to the development of antiviral nucleotides is the off-target effects, caused primarily by cellular polymerase utilization of the compound^7^. However, the specificity of antiviral nucleotides continues to improve^8^.

Recently, a new class of antiviral ribonucleotide was approved in Japan to treat influenza virus infection^9^. The drug has been named favipiravir, and is also known as T-705. Favipiravir is a fluorinated base analogue with a pyrazine carboxamide (**Supplementary Fig. 1A**). This compound requires the cellular nucleotide salvage pathway to convert the base into a nucleoside triphosphate. A version of favipiravir lacking fluorine, known as T-1105 (**Supplementary Fig. 1B**), is also active, but its conversion to the triphosphate is less efficient^10^. The nucleoside version of T-1105 is referred to as T-1106 (**Supplementary Fig. 1C**), and the metabolism of this compound to the triphosphate yields a drug with efficacy superior to favipiravir^10^. Studies of the mechanism of action of these compounds have resulted in ambiguity. Some studies were consistent with chain termination^11^, while others were consistent with lethal mutagenesis^12^. Of course, such uncertainty in the mechanism of action complicates approval beyond Japan.

In previous work, we used a high-throughput magnetic-tweezers platform to monitor hundreds of individual poliovirus (PV) RdRp elongation complexes over thousands of nucleotide-addition cycles and observed the ability of T-1106 to cause the RdRp to pause and then backtrack^13^. The elongation complex could recover from the backtracked state, but recovery required tens to thousands of seconds. As a result, traditional polymerase elongation assays would view these backtracked states as prematurely terminated products^13^. This backtracking phenomenon was not observed with prototypical chain terminators or mutagens. These studies therefore provided very compelling evidence for the existence of a third mechanistic class of antiviral ribonucleotide analogs.

While the structure of the backtracked state is not known, the nascent RNA was likely displaced from template to yield a single-stranded 3’ end with lengths greater than several tens of nucleotides^13^. The ability of an RdRp to produce free single-stranded ends was intriguing, because such an end could undergo an intermolecular template switch by annealing to a second template, with resumed synthesis producing a recombinant RNA product. The goal of the present study was to determine the extent to which the backtracked state represented an intermediate on path for recombination. To do so, we introduced the RdRp from EV-A71, a virus known be more prone to recombination than PV^14^, into our pipeline with the idea that perhaps such backtracked states might be more prevalent with this enzyme.

By comparing the EV-A71 and PV RdRps, we provide evidence that the backtracked state is indeed an intermediate for template switching. Unexpectedly, for EV-A71 RdRp we observed a high frequency of intramolecular template switching, which has been coined “copy-back RNA synthesis” in the literature^15^. Our data for EV-A71 RdRp also indicate that the pyrazine-carboxamide class of antiviral ribonucleotides enhances such switching and may function by promoting the formation of defective viral genomes^15^. Our study therefore makes a clear mechanistic connection between recombination and copy-back RNA synthesis, and provides compelling evidence that an enhancement of the probability of template switching results in an antiviral effect. The experimental paradigm reported here should prove useful in dissecting the contributions of the RdRp and the template capable of promoting formation of defective viral genomes, a newly emerging strategy for interfering with virus multiplication and attenuating viral pathogenesis^15^.

## RESULTS

### Pausing of EV-A71 RdRp promotes copy-back RNA synthesis

The development of a magnetic-tweezers platform to monitor nucleotide addition by nucleic acid polymerases is unmasking the stochastic behavior of viral polymerases and illuminating states of the polymerase induced by pausing and drugs^13,16,17^. In this assay (Fig. 1A), single-stranded RNA is tethered to the surface and a magnetic bead. A template ssRNA, including a 24 nt hairpin structure at its 3’ end, is annealed to the tethered ssRNA strand, creating a predominantly double-stranded RNA. Primer-extension from the 3’-end of template RNA will lead to displacement of the template RNA from the tethered RNA. At forces > 8 pN, the conversion of dsRNA to ssRNA causes a corresponding increase in the distance of the bead from the surface, thus permitting measurement of nucleotide incorporation with few nucleotide resolution over thousands of cycles of nucleotide addition^13,16,17^.

**Figure 1.**
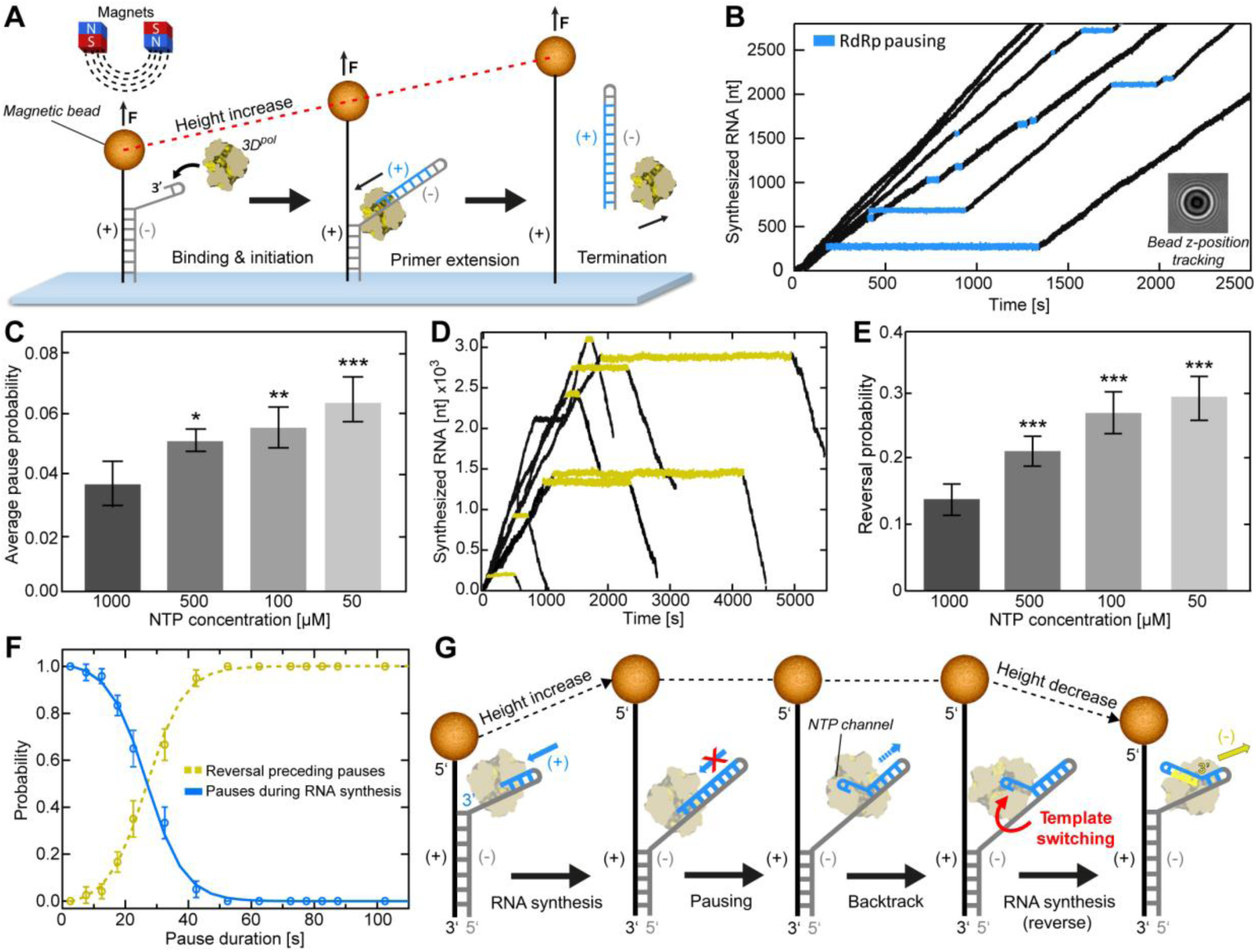
Magnetic tweezers assay of EV-A71 RdRp reveals pause-dependent copy-back RNA synthesis. (**A**) Schematic of the single-molecule (+)-strand RNA synthesis assay, showing the binding of an RdRp to a hairpin at the 3’-end of the (-)-strand of the surface-attached RNA template. A magnetic bead attached to the RNA construct is subject to a constant force of 25 pN. (**B**) Sample EV-A71 RdRp trajectories showing stochastic pausing behavior over time, inferred *via* the change of the diffraction pattern of the attached magnetic bead (inset) at a rate of 50 Hz. Within individual trajectories, the data in blue highlight single pauses of different duration and template position during processive RNA synthesis. (**C**) Comparison of the average pause frequency (±SD) at different rNTP concentrations extracted from dwell-time distributions (**Supplementary Fig. 2A**). Dataset statistics and experimental parameters for each condition we provided in **Supplementary Table 1**. (**D**) Sample individual RdRp elongation trajectories showing the occurrence of reversal events following extended pausing (yellow). (**E**) The average occurrence probability (±SD) of reversals per RdRp for all measured conditions. (**F**) The probability of observing pauses of specified pause durations during RNA synthesis (blue) co-plotted with the probability of reversals (yellow). (**G**) Proposed model underlying the reversal events, in which pause-induced elongation complex is followed by copy-back RNA synthesis of the newly synthesized (+)-strand RNA (blue). Synthesis of a new (-)-strand (yellow) results in reannealing of the original (+)-strand template (grey) to the complementary ssRNA tether (black), leading to an overall decrease in RNA tether extension. Statistical analyses were performed using one-way analysis of variance (ANOVA) with comparative Tukey post-hoc test (significance levels α: *** = 0.001; ** = 0.01; * = 0.05). See also **Supplementary Fig. 2**.

In a previous study with PV RdRp, we showed that one consequence of prolonged pausing by the enzyme is backtracking, where the enzyme unwinds the 3’-terminus of nascent RNA^13^. Such a state had not previously been reported, but we envisioned that it could serve as an intermediate for homologous recombination by an intermolecular template-switching mechanism. Such a template switch would occur by annealing of the 3’-terminus of the backtracked (donor) template to a second (acceptor) template. In the magnetic tweezers, there is no acceptor template; therefore, intermolecular template switching does not occur. In most instances, the nascent RNA reannealed to the original template and RNA synthesis resumed^13^.

While recombination clearly occurs in PV, the frequency of recombination appears higher in EV-A71^14^. The expectation therefore is that EV-A71 RdRp might exhibit the backtracked state at a higher frequency. As illustrated in Fig. 1B, processive nucleotide addition by EV-A71 RdRp (quantified in **Supplementary Fig. 2C**) was frequently interrupted by pauses of differing duration. For PV RdRp, we previously observed an inverse correlation between nucleotide concentration and pause probability^13^. EV-A71 behaved similarly (Fig. 1C and **Supplementary Fig. 2A,B**). Pausing is thought to occur in response to misincorporation and happens more often at low nucleotide concentration^13^. Such a response might facilitate correction by pyrophosphorolysis^13,18^.

At some frequency for PV RdRp, however, the paused state undergoes backtracking^13^. Surprisingly, under the same conditions, EV-A71 RdRp can undergo an apparent reverse translocation (referred to as “reversals” in what follows), as the position of the bead is observed to decrease relative to the surface (Fig. 1D). Pauses induced by low nucleotide concentrations, found to be drivers of backtracking by PV RdRp^13^, are also drivers of reversals by EV-A71 RdRp (Fig. 1E,F). While it was not possible to identify sequence motifs that trigger reversals, G:C-rich regions exhibited a significantly higher probability of reversals (**Supplementary Fig. 3**). Given the assay design, the only way to decrease the position of the bead relative to the surface would be to reanneal the displaced template ssRNA to the tethered ssRNA. For this to happen, nascent ssRNA would have to be displaced from the template ssRNA. One probable explanation for this observation is that pausing by EV-A71 RdRp leads to backtracking, which produces a single-stranded 3’-end that is used by EV-A71 RdRp as a primer for copy-back RNA synthesis (Fig. 1G). The use of a single polymerase for the initial round of RNA synthesis and copy-back RNA synthesis is supported by the good agreement of the kinetic behavior of the polymerases during both processes (**Supplementary Fig. 2D**).

### A recombination-deficient EV-A71 RdRp variant attenuates virus population

Our studies make a compelling case for the magnetic tweezers providing a glimpse into intermediates leading to intermolecular template switching (homologous recombination) and copy-back RNA synthesis (intramolecular template switching). The availability of a recombination-defective EV-A71 RdRp derivative would prove useful in making the strongest case for a relationship between the observation of backtracking or reversals and recombination.

The Andino and Barton laboratories identified Y275H in PV RdRp as a substitution that impaired recombination substantially, among several others^19,20^. The molecular basis for the reduced recombination efficiency is not known^19,20^. The orthologous substitution is Y276H in EV-A71 RdRp. We engineered the Y276H substitution into RdRp-coding sequence of EV-A71. In the context of a subgenomic replicon, Y276H EV-A71 replicated better than WT (Fig. 2A). By plaque assay and RT-qPCR, Y276H and WT EV-A71 were indistinguishable (Figs. 2B,C). To assess the recombination efficiency, we transfected the subgenomic replicon RNA described above (donor template) and an EV-A71 genomic RNA deleted for 3D^pol^-coding sequence (acceptor template) into cells; recombination between these RNAs produces viable virus (Fig. 2D). The yield of recombinant virus was reduced by nearly 100-fold (Fig. 2E).

**Figure 2.**
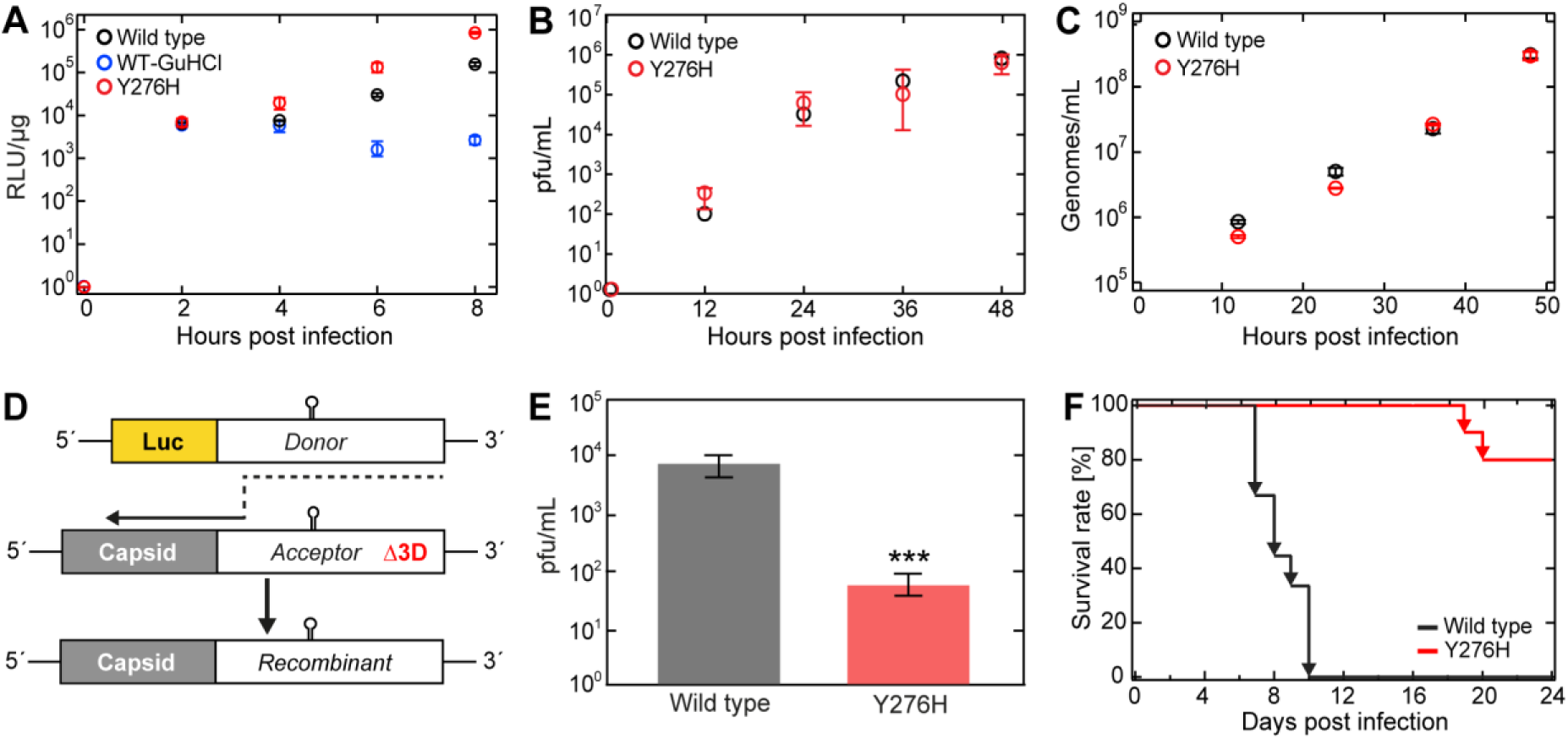
EV-A71 Y276H RdRp variant exhibits significance decrease in recombination efficiency and virulence. (**A**) Replication efficiency of the EV-A71 wild type and Y276H variant subgenomic replicons in RD cells as a function of time post transfection. GuHCl refers to the presence of guanidine hydrochloride, a potent inhibitor of replication. Luciferase activity is reported in relative light units (RLU; AVG ±SD) per microgram of total protein (n = 3 replicates per time point). (**B**) EV-A71 wild type (black) and Y276H variant (red) single-step growth curves (±SD; n *=* 3) confirm similar plaque formation. (**C**) EV-A71 wild type (black) and Y276H variant (red) single-step growth curves (AVG ±SD; n = 3 for each time point). RD cells were infected with virus equivalent to 200 genomes/cell. Samples were taken at the indicated times and the genome amount of virus titer was quantified via RT-qPCR. (**D**) Cell-based recombination assay. EV-A71 C2-strain firefly luciferase-encoding sub-genomic replicon (donor) and full length EV-A71 C2-MP4 strain genome (acceptor) carrying a lethal deletion of the 3D^pol^ region are co-transfected in RD cells. Only upon co-transfection, replication-competent virus can be generated by RdRp-mediated template switch from donor to acceptor (indicated by dashed black arrow). (**E**) The Y276H mutation in the EV-A71 replicon inhibits recombinant yield. Resulting recombinant virus were quantified by pfu/ml (AVG ±SD; *n = 3*). (**F**) EV-A71 C2-MP4 wild type and Y276H virulence in hSCARB2 mice. 21-day old mice were orally inoculated with either wild type or Y276H virus (n = 10 per virus) at a dose equivalent to 2×10^7^ genomes and scored for survival post-infection. Effect of Y276H variant is severely attenuated relative to that of WT. Statistical analysis consisted of an unpaired, two-tailed t-test (significance level P: *** ≤ 0.001)

Previous studies have indicated that the ability of PV RdRp to catalyze recombination is required for PV to cause disease in a mouse model, even though growth in cell culture appears equivalent to or better than WT. We evaluated the virulence of Y276H EV-A71 in a mouse model that supports infection via oral inoculation. These mice express human SCARB2 protein, which is a receptor for EV-A71^21^. The population of viruses carrying the Y276H EV-A71 was highly attenuated in this model relative to WT (Fig. 2F), in agreement with a recently published study^22^.

### Recombination deficiency of Y275(6)H RdRps is caused by enhanced binding to nascent RNA and diminished backtracking

Based on the data gathered in the past for PV RdRp and that described above for EV-A71 RdRp, our mechanism for template switching begins with RdRp pausing, followed by backtracking, and ending with either resolution of the backtrack (PV) or reversals (EV-A71). The availability of a recombination-defective EV-A71 RdRp, Y276 RdRp, should facilitate establishment of a correlation between events observed using the magnetic tweezers and requirements of the template-switching process.

The kinetics of correct nucleotide addition were unchanged for Y276H RdRp relative to WT (note overlap of probability density in the dwell-time distributions up to 3 s in Fig. 3A). However, the rate of processive nucleotide addition for the recombination-defective derivative appeared compromised by a 2-fold increase in pausing frequency (Fig. 3B) and a 4-fold increase in the duration of the pause (Fig. 3C). The overall processivity of this derivative was reduced by 4-fold (Fig. 3D). In spite of the finding that pausing was increased substantially, reversals were actually reduced by 4-fold (Fig. 3E).

**Figure 3.**
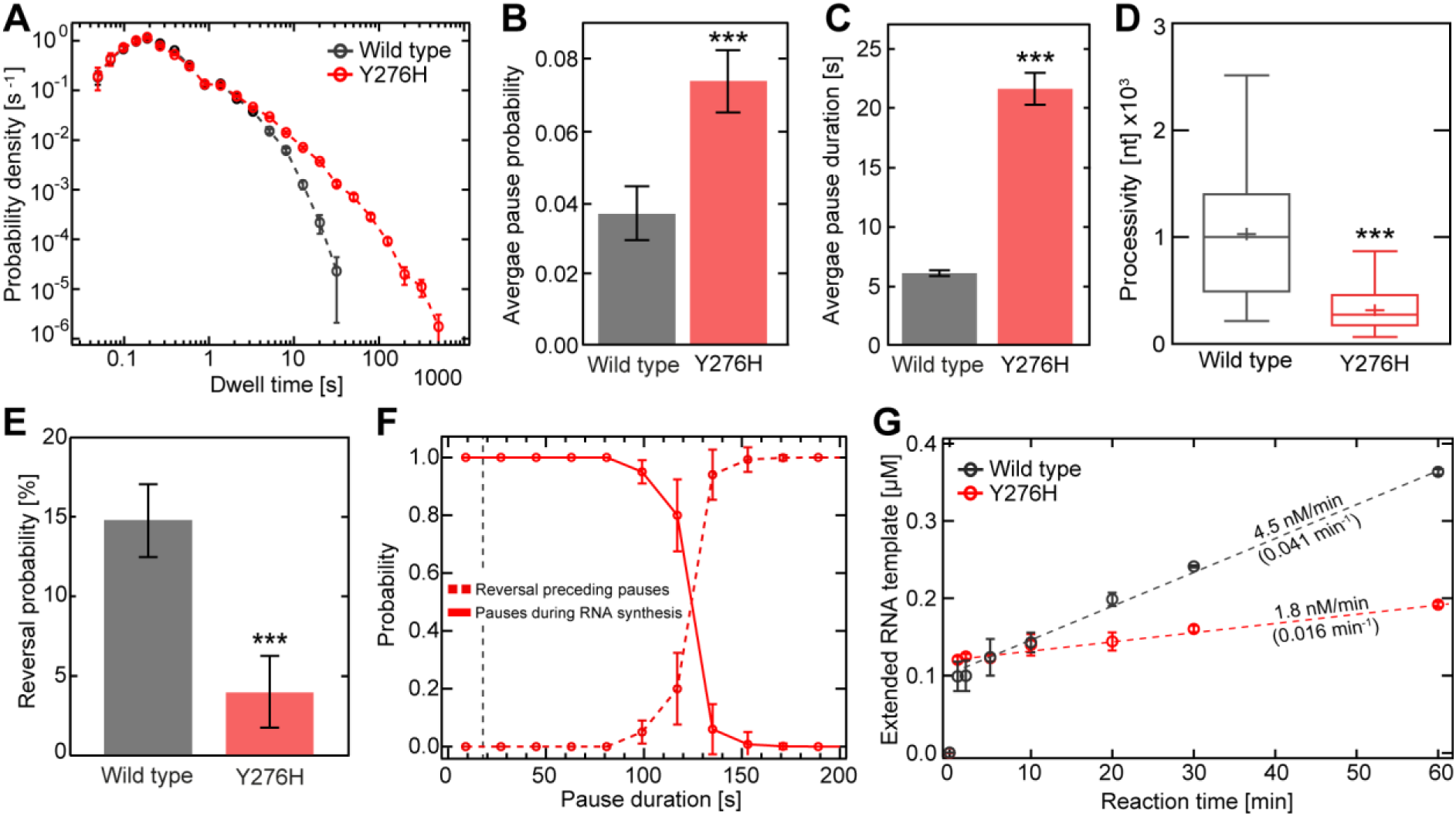
EV-A71 Y276H RdRp variant is impaired for copy-back RNA synthesis. (**A**) Superimposed dwell-time distributions for EV-A71 Y276H RdRp (red) and WT (black) RdRp. The dwell times used for the construction of the distributions are the time needed for the RdRps to synthesize four consecutive nucleotides. The Y276H variant exhibits a broad increase in the probability and duration of long pauses compared to wild type. The error bars (AVG ±SD) result from bootstrapping with 1,000 iterations. (**B-D**) Quantification of the data in panel (A). In comparison to WT RdRp, the Y276H variant shows a significant increase in (**B**) average pausing probability (±SD) and (**C**) average apparent pause duration (±SEM). The processivity (**D**) is significantly decreased for the Y267H RdRp variant compared to WT. (**E**) The Y276H RdRp (red) variant causes significantly fewer reversal events than WT RdRp (grey). (**F**) The probability of observing pauses of specified durations during RNA synthesis (red line) co-plotted with the probability of observing reversals (red dashed line). (**G**) *In vitro* bulk RNA synthesis assay results (AVG ±SD; n = 3 repetitions) showing the amount of extended Sym/SubU template for EV-A71 wild type and Y276H RdRp over time. Dashed lines represent linear regressions fitted to the data, revealing significantly lower approximated rates of RNA extension (values above the dashed lines) and turnover (values below the dashed lines; unit: RNA per minute per RdRp) for the Y276H RdRp variant compared to WT. Statistical analyses were performed using unpaired, two-tailed t-tests (significance level P: *** ≤ 0.001).

The reversals observed with Y276H RdRp derived from a pause-dependent mechanism as observed for WT (Fig. 3F). With pausing elevated and reversals diminished, the backtracked intermediate should accumulate. This was not the case, suggesting that an inability to backtrack is the molecular defect associated with the Y276H RdRp derivative that interferes with recombination.

For backtracking to occur, the paused polymerase needs to release the 3’-end. The stability of the RdRp at the 3’-end can be inferred from the steady-state rate constant for single-nucleotide incorporation^23^. A reduction in the value of this rate constant means increased affinity for the terminus. Y276H RdRp elongation complex was 3-fold more stable than that formed by WT on the synthetic template developed to monitor elongation by enteroviral RdRps (Fig. 3G)^23^. It is likely that the increased frequency of pausing and longer duration of the pause are caused by enhanced affinity of the Y276H RdRp for the 3’-end of nascent RNA.

The availability of the orthologous derivative in PV, Y275H RdRp, provided the opportunity to determine if the mechanism causing the recombination defect is the same for both viral polymerases. For PV RdRp, a decrease in backtracking should be discernible because backtracking is its end point, providing empirical validation of our inference above. As shown in Fig. 4A-D, Y275H RdRp behaved as Y276H RdRp (based on features of the dwell-time distribution, including pause probability and average pause duration, as well as processivity) relative to its WT RdRp. Importantly, however, the probability of backtracking was reduced for Y275H RdRp relative to WT RdRp (Fig. 4E). For PV RdRp, backtracking derived from a pause-dependent mechanism (Fig. 4F), as the reversals did for EV-A71 RdRp (Fig. 1F). The significant reduction in backtracking probability for Y275H RdRp may reduce the incidence of template-switching-proficient backtracks. As for the recombination-defective EV-A71 polymerase, Y275H RdRp exhibits a substantially more stable elongation complex than WT (Fig. 4G).

**Figure 4.**
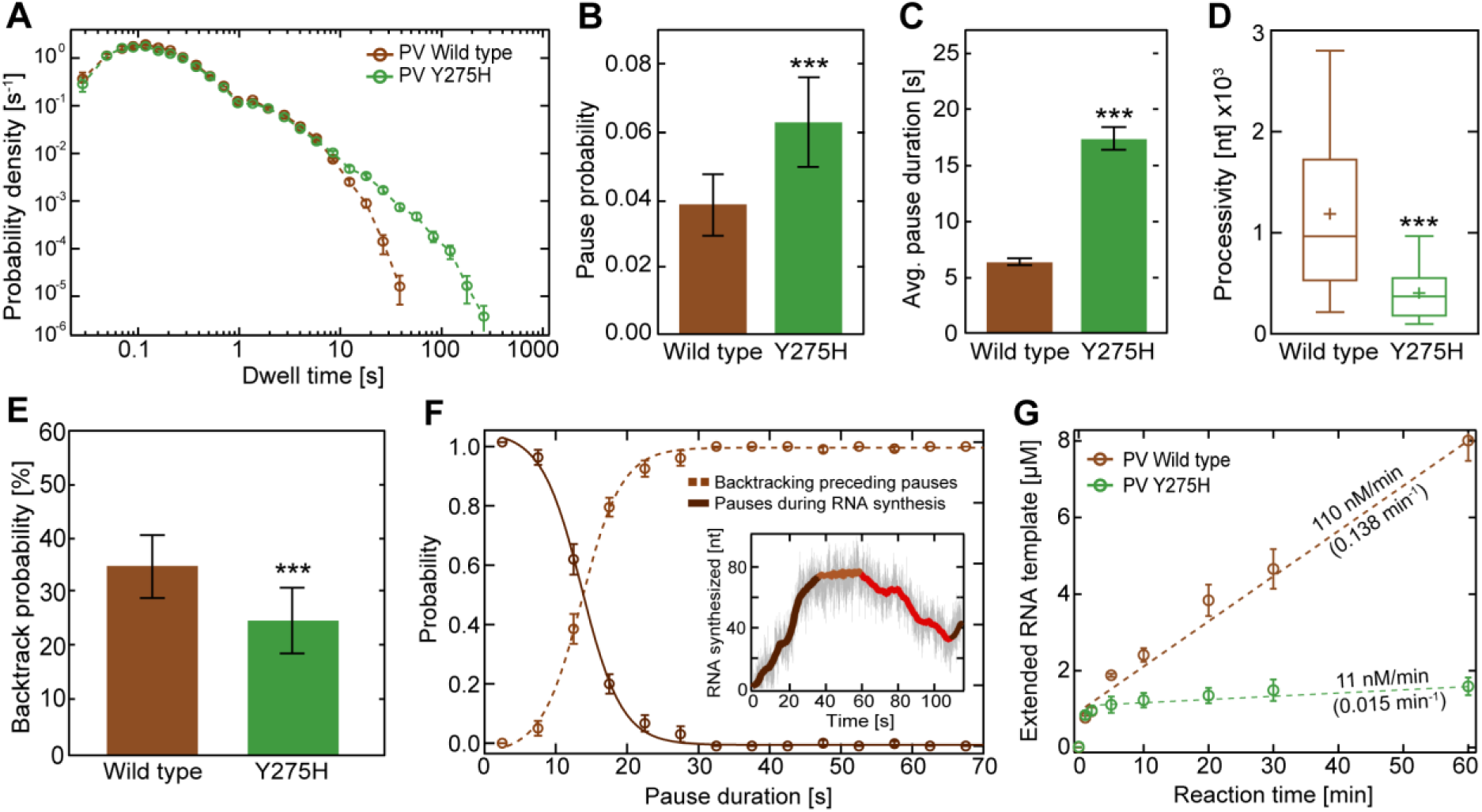
PV Y275H RdRp is impaired for backtracking. (**A**) Superimposed dwell-time distributions of poliovirus Y275H RdRp variant (green) and WT (brown) RdRp. The 275H variant exhibits a broad increase in the probability and duration of long pauses compared to wild type. The dwell-time window was set to 4 nt, and the error bars (AVG ±SD) result from bootstrapping with 1,000 iterations. (**B-D**) Quantification of the data in panel (A). In comparison to WT RdRp, the Y275H variant shows a significant increase in (**B**) average pausing probability (±SD) and (**C**) average apparent pause duration (±SEM) during RNA synthesis. The processivity (**D**) is significantly decreased for the Y265H RdRp variant compared to WT. (**E**) The Y275H variant (green) shows a significantly decreased backtracking probability (F, inset) than WT RdRp (brown). (**F**) The probability of observing pauses of specified durations during PV WT RNA synthesis (dark brown) co-plotted with the probability of observing backtracking (brown dashed line). (**G**) *In vitro* bulk RNA synthesis assay results (AVG ±SD; n = 3 repetitions) showing the amount of extended Sym/SubU template for PV WT and Y275H RdRp over time. Dashed lines represent linear regressions fitted to the data, revealing significantly lower approximated rates of RNA extension (values above the dashed lines) and turnover (values below the dashed lines; unit: RNA per minute per RdRp) for the Y275H RdRp variant compared to WT. Statistical analyses were performed using unpaired, two-tailed t-tests (significance level P: *** ≤ 0.001).

### Conformational dynamics of the RNA-binding channel as a determinant of the type and efficiency of RdRp-catalyzed template switching

It has become increasingly clear that the biochemical properties of enzymes are governed as much by their conformational dynamics as by their three-dimensional structure^24,25^. The conformational dynamics of PV RdRp is an important contributor to the specificity and efficiency of its polymerase function^26^. The availability of structures of the RdRp from PV and EV-A71^27,28^ and models of the Y276H RdRp produced for this study, permitted us to determine whether structure and/or dynamics explain the differences observed in the propensity for one RdRp to catalyze copy-back RNA synthesis but the other being incapable of this activity or explain the impact of the Y275(6)H substitution on backtracking or intramolecular template switching. Structural differences provided little, if any, insight (Fig. 5A,B); therefore, we turned to dynamics.

**Figure 5.**
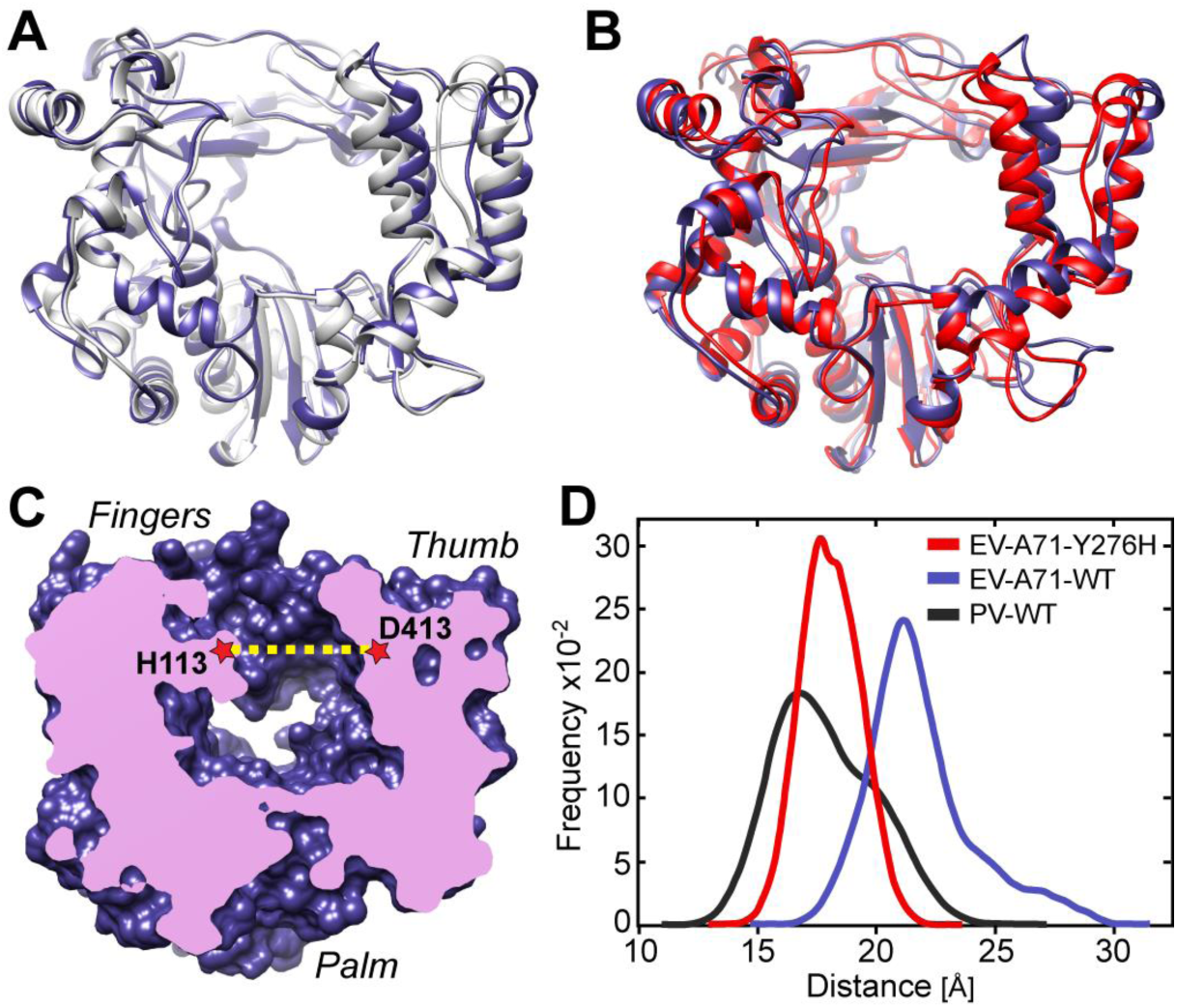
RNA-duplex channel dimensions and conformational dynamics of EV-A71 and PV RdRps. (**A**) Superimposed crystal structures of EV-A71 WT (blue) and PV WT (grey) shown as cartoons. (**B**) Superimposed crystal structures of EV-A71 WT (blue) and major conformation of its Y276H mutant (red) that result from MD simulations. (**C**) Cut-through volume rendering of EV-A71 WT crystal structure (PDB 3N6L), where the RNA duplex channel can be observed in the center of the structure. The channel width was assessed by measuring the distance between His-113 at the fingers and Asp-413 at the thumb domains in EV-A71, or their equivalent residues Ser-112 and Asp-412 in the PV WT crystal structure (PDB 1RA6). (**D**) RNA duplex channel widths measured for EV-A71 WT (blue) and PV WT (black) from their crystal structures, and from MD simulations for the EV-A71 Y276H variant (red).

We have established a pipeline for evaluating the conformational dynamics of picornaviral polymerases by using molecular dynamics simulations (MD)^25,29^. Our analysis evaluates correlated motions of the conserved structural motifs, and in particular reports on the dynamics of the open and occluded states of the catalytic site that drive specificity of nucleotide selection^29^. We also evaluate dynamics of the RNA-binding channel. One approach is to monitor the time-dependent changes in the distance of two residues bordering the channel: H113 and D413 in case of EV-A71 (Fig. 5C)^29^. This latter analysis was most informative, since it showed that the average size of the RNA-binding channel of PV RdRp was smaller (by ~4 Å) than that of EV-A71 RdRp (Fig. 5D). Interestingly, introduction of the Y7276H substitution decreased the average size of the RNA-binding channel (by ~3 Å) over the duration of time sampled (Fig. 5D).

### RdRp backtracking in response to incorporation of the pyrazine carboxamide, T-1106, promotes intra- and intermolecular template switching

Our study thus far makes a very compelling case for the ssRNA 3’-end produced by RdRp backtracking serving as an intermediate for both intra- and intermolecular template switching. Our interest in backtracking, however, was motivated by our previous observation that incorporation of T-1106 ribonucleoside triphosphate (T-1106-TP) into nascent RNA induced backtracking^13^. In the context of our current findings, we would expect T-1106 ribonucleotide to promote copy-back RNA synthesis and/or homologous recombination. If this is the case, then T-1106 may actually represent the first antiviral ribonucleotide whose antiviral activity can be attributed to a post-incorporation event unrelated to termination or mispairing, in this case template switching.

In the first series of experiments, we compared EV-A71 RdRp elongation dynamics in the magnetic tweezers in the absence and presence of T-1106-TP. Incorporation of T-1106 ribonucleotide induced elevated levels of pausing (Figs. 6A-C; **Supplementary Fig. 4A**), diminished processivity (**Supplementary Fig. 4B**), and an increased incidence of reversals (Fig. 6D), reflecting increased intramolecular template switching (copy-back RNA synthesis). We also verified that the outcome of T-1106-TP utilization on PV RdRp was as expected based on our previous study (**Supplementary Fig. 5**^13^. T-1106 induced backtracking by PV RdRp (**Supplementary Fig. 5D**), consistent with a greater likelihood for intermolecular template switching (homologous RNA recombination).

**Figure 6.**
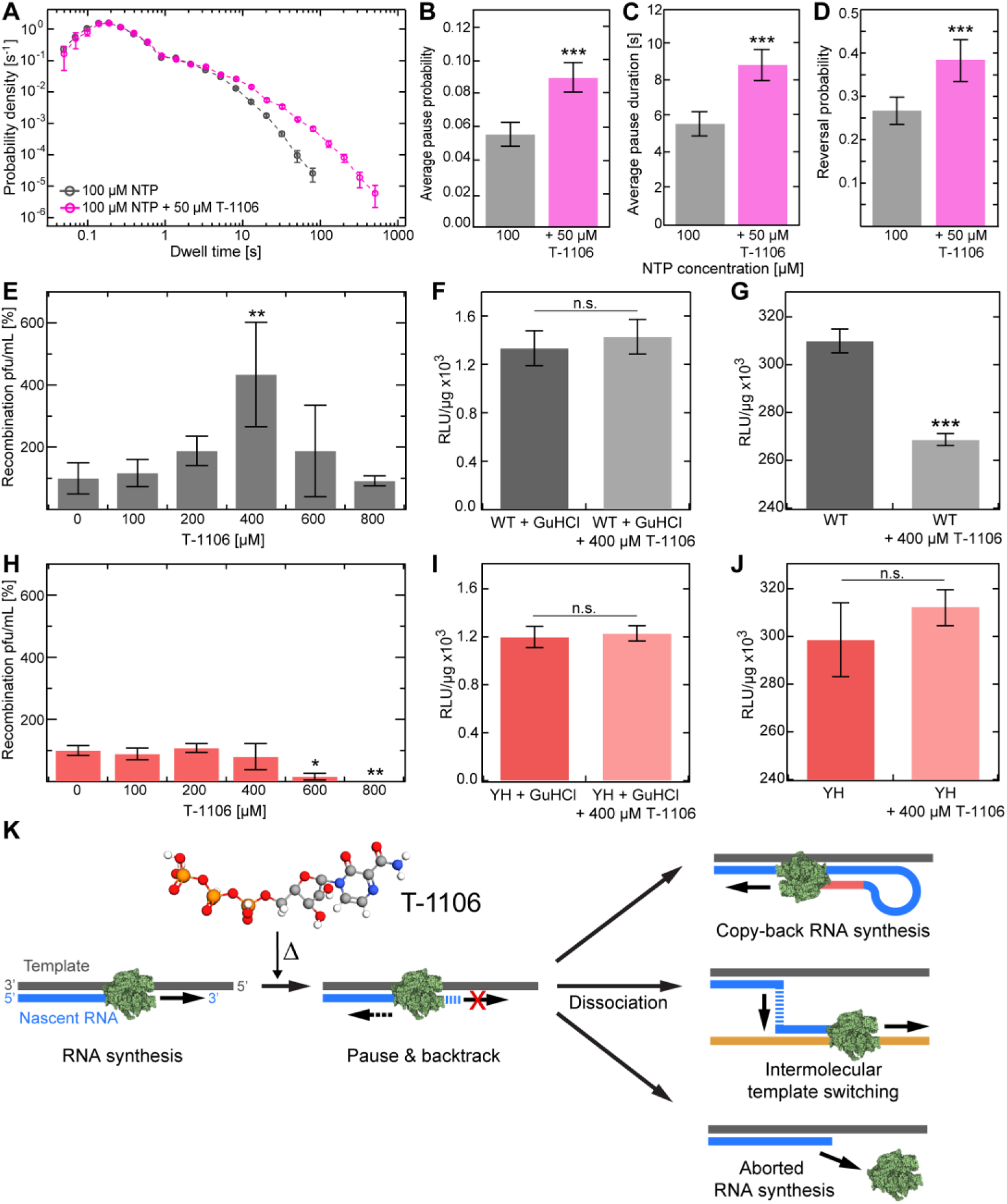
Pyrazine carboxymide T-1106 induces an increase in intra- and intermolecular template switching. (**A**) Superimposed dwell-time distributions of EV-A71 RdRp RNA synthesis activity in the presence (magenta) or absence (grey) of the nucleotide analogue T-1106. The dwell-time window was set to 4 nt, and the error bars (±SD) result from bootstrapping with 1,000 iterations. (**B-D**) The addition of T-1106 changes significantly the pausing behavior of wild type RdRp, exhibiting significantly increased (**B**) pausing probability (±SD) and average pause duration (±SEM), leading to (**D**) an increase in copy-back RNA synthesis probability. (**E**) Cell-based recombination assays conducted in presence of different concentrations of T-1106. Relative viable wild type recombinant yield, normalized as a percentage of a carrier (DMSO) treated control (AVG ±SD; n = 3 replicates for each condition). (**F**, **G**) EV-A71 donor translation (**F**) and replication (**G**) efficiency (AVG ±SD) for wild type RdRp. (**H**) Relative Y276H viable recombinant yield normalized as a percentage of a carrier (DMSO)-treated control (AVG ±SD; n = 3 for each condition). (**I**, **J**) EV-A71 donor translation (**I**) and replication (**J**) efficiency (AVG ±SD) for Y276H RdRp variant. (**K**) Hypothesized mechanism of antiviral-caused stall in RNA synthesis leading to copy-back RNA synthesis, intermolecular template switching, or abortive genome synthesis by RdRp dissociation. Statistical analyses in (E, H) were performed using one-way analysis of variance (ANOVA) with comparative Tukey post-hoc test (significance levels α: *** = 0.001; ** = 0.01; * = 0.05), and unpaired, two-tailed t-tests (significance level P: *** ≤ 0.001). See also **Supplementary Figs. 4**, **5** and **6**.

We recently adapted a cell-based assay for PV RNA recombination to EV-A71 ^14^, thus providing an approach to evaluate the impact of T-1106 on EV-A71 RNA recombination in cells. T-1106 inhibits EV-A71 multiplication (**Supplementary Fig. 6A**). However, between 200 µM and 600 µM T-1106 a clear increase in recombination was observed (Fig. 6E). Using a subgenomic replicon expressing luciferase, we showed that T-1106 did not affect reporter expression in the presence of the replication inhibitor guanidine hydrochloride (GuHCl), consistent with the drug having no effect on translation of the viral polyprotein (Fig. 6F). Replication-dependent, reporter expression was inhibited (Fig. 6G). Antiviral ribonucleotides that function by lethal mutagenesis do not significantly reduce replication-dependent, reporter expression^28^. It is therefore possible that the observed inhibition reflects a reduction in the amount of replication-competent replicon RNA resulting from enhanced copy-back RNA synthesis in the presence of T-1106 and production of truncated (defective) replicons.

The studies presented above are consistent with T-1106 promoting copy-back RNA synthesis and homologous RNA recombination. We performed the same experiments with the recombination-defective variant, Y276H EV-A71. With this variant, T-1106 failed to increase recombination (Fig. 6H), consistent with the RdRp serving as the mediator of the effect of T-1106. Using the Y276H-encoding subgenomic replicon, we showed that translation was not impacted by the presence of T-1106, as expected (Fig. 6I). Unexpectedly, replication-dependent reporter expression was also not impaired by the presence of T-1106 (Fig. 6J). This observation confirms that the reduction in replication-dependent reporter expression was indeed caused by a backtracking-induced phenomenon. Of the few possibilities, defective replicons produced by copy-back RNA synthesis is a likely possibility^15^. It was not possible to evaluate production of the defective genomes directly because of the rapid degradation of subgenomic replicon RNA that occurs immediately following transfection.

We have discovered that the T-1106-induced backtracking/reversals observed *in vitro* actually correspond to the induction of homologous RNA recombination and/or copy-back RNA synthesis in cells. The impact of the drug on these template switching-dependent mechanisms is alleviated for the Y275(6)H, recombination-defective variant. Does this mean that a facile route to the development of resistant to this mechanistic class of antiviral ribonucleotides exist? To address this possibility, we evaluated the sensitivity of Y276H EV-A71 to T-1106. There was no more than a 2-fold change in the observed sensitivity of this derivative to T-1106 relative to WT (compare **Supplementary Fig. 6B** to **Supplementary Fig. 6A**). Therefore, elimination of the template-switching dependent activity is insufficient for resistance to T-1106 and perhaps all members of the pyrazine-carboxamide class of antiviral ribonucleotides.

## DISCUSSION

Among the last frontiers in RdRp enzymology is a mechanistic description of RNA recombination. In the majority of RNA viruses, recombination is primarily an RdRp-mediated process. In PV, which is one of the most studied RNA virus models, it has been that recombination involves an RdRp-mediated template-switching mechanism in cells^30^ and that RdRp is sufficient to catalyze a template-switching reaction *in vitro*^31^. Over the past few years, a renaissance in the study of RNA recombination has begun, as evidenced by the development of cell-based assays to probe the incidence of recombination^14,32,33^.

In our earlier work that monitored PV RdRp elongation dynamics at the single-molecule level *in vitro*, we observed extensive RdRp pausing followed by backtracking in response to incorporation of the pyrazine carboxamide ribonucleotide analogue T-1106^13^. In a typical single-molecule experiment, we observed the production of tens of nucleotides ssRNA caused by PV RdRp backtracking, which significantly increases in probability in response to incorporation of T-1106^13^, and here further quantified in **Supplementary Fig. 5E**). Given sufficient time, the drug-induced ssRNA reannealed, the RdRp rebound to the primer-template junction, and elongation resumed.

Observation of such spatiotemporal dynamics of the nascent RNA-template-RdRp complex was unprecedented and motivated further study. Intriguingly, cell-based recombination studies in PV and other RNA viruses have shown that T-1106 significantly increased recombination efficacy (Fig. 6E)^34–38^. Since PV RdRp backtracking propensity represented the sole change in RNA synthesis dynamics in response to T-1106 treatment, it could represent a recombination intermediate. With that possibility in mind, this study added the EV-A71 RdRp, because EV-A71 has been documented to undergo substantially more recombination in nature than observed for PV^14^. EV-A71 recombination is a major cause of the recurring outbreaks in Asia^39^.

The first major conclusion of this study is that the viral RdRp has evolved to sense and to respond to incorporation of an incorrect nucleotide or nucleotide analogue by pausing and backtracking, thereby producing a recombinogenic 3’-end. Nucleotide misincorporation will be increased at lower nucleotide concentrations and in the presence of skewed nucleotide pools^13^. Under these conditions, both EV-A71 and PV RdRps exhibit an increase in the probability and duration of pausing (Fig. 1C and **Supplementary Fig. 2B**, respectively)^13^. Similarly, utilization of T-1106 triphosphate increases the pause probability and duration for both polymerases (EV-A71: Fig. 6B,C and **Supplementary Fig. 2B**; PV: **Supplementary Fig**. 5**B,C**). The average pause duration is measured in the tens of seconds for both enzymes (EV-A71: **Supplementary Fig. 2B**; PV: **Supplementary Fig**. 5**C**), not milliseconds as typically observed for nucleotide addition^40^. We suggest that the consequence of a misincorporation- or analogue-induced pause is backtracking in both systems. In the case of PV RdRp, there is no debate because the backtracked state was observed to accumulate during the course of an experiment (Fig. 4E) and does so more frequently upon utilization of T-1106 triphosphate (**Supplementary Fig. 5E**)^13^. With EV-A71 RdRp, the backtracked state does not accumulate. Instead, the probability of apparent RdRp backward translocation increases (Figs. 1D, 6D), which reflects nothing more than the use of the 3’-end of nascent RNA as a primer for copy-back RNA synthesis (Fig. 1G). Backtracking would be the simplest mechanism for liberating the 3’-end for use as a primer.

While there has been some speculation that RNA recombination and copy-back RNA synthesis are mechanistically related based on observations that circumstances that elevate RNA recombination also elevate copy-back RNA synthesis^15^, how the two processes are connected is not known. Our study is consistent with two fates for the ssRNA produced by backtracking. The first is intermolecular template switching, which is homologous RNA recombination. The second is intramolecular template switching, which is copy-back RNA synthesis. Therefore, this study makes an unambiguous mechanistic link between these two processes by showing that they share the same backtracking intermediate, which is produced by the same triggers.

Why are there two different outcomes for the backtracked state, which does not accumulate with the EV-A71 RdRp, but does with PV RdRp? For copy-back RNA synthesis to occur, the nucleic-acid-binding site needs to be sufficiently large to accommodate a three-stranded intermediate at the time of initiation (Fig. 1G). Analysis of the dynamics of the nucleic-acid-binding site of EV-A71 RdRp demonstrated a clear ability of this enzyme to accommodate such an intermediate with an approximate diameter of ~24Å. (Fig. 5D). The corresponding site in PV RdRp is on average 4 Å smaller (Fig. 5D). Perhaps, the dimensions of PV RdRp are too small to accommodate such an intermediate. If this is the case, then the dimensions of the RNA-binding site sampled by an RdRp may predict the capacity of the polymerase to catalyze copy-back RNA synthesis.

The product of copy-back RNA synthesis will likely be a truncated dsRNA that is defective in its coding capacity and/or a potent activator of innate immune responses^15^. Whether or not production of copy-back RNA is deliberate is unclear but may be for some viruses because the copy-back RNA can dampen the intensity of the infection^15^. During replication in the cell when the backtracked state arises, templates complementary to the single-stranded 3’-end would be present. Under these conditions, perhaps intermolecular template switching would perhaps be favored for both enzymes.

Over the past few years, a connection has been made between RdRp fidelity and recombination^14,32,39,41,42^. With more incorporation errors and/or consumption of nucleotide analogues by the RdRp comes an increase in the frequency of recombination both in cells and in test tubes^39^. The reason for this has not been clear. Our studies provide a mechanism. Recombination is thought to provide a means to suppress the impact of deleterious mutations^40,43,44^. Our data would suggest that a balance between mutation and recombination must exist, because increasing the frequency of recombination exhibits antiviral activity (Fig. 6E, **Supplementary Fig. 6A**) attributable to production of defective genomes (Fig. 6G). Consistent with too much recombination being inhibitory to virus genome replication, not every event that could trigger recombination (Figs. 1C, 6B, and **Supplementary Fig. 5B**) actually does so; only a fraction of paused polymerases actually undergo copy-back synthesis (Figs. 1E, 6D) or accumulate in the backtracked state (**Supplementary Fig. 5E**). Our previous studies of PV RdRp suggested that only a subset of elongation complexes was competent for misincorporation or incorporation of certain nucleotide analogues^13^. If this is the case, then these misincorporation-competent elongation complexes may be the only complexes competent for backtracking and template switching.

For an event to trigger template switching, that event must first trigger backtracking. Our studies have focused on the perception of a nucleotide addition as correct or incorrect as a mechanism to induce backtracking and consequentially template switching. However, other mechanisms to induce backtracking may exist. One intriguing possibility is that certain template sequences may direct the RdRp to pause and/or backtrack. Sequence- and factor-dependent pausing is well represented in the DNA-dependent RNA polymerase literature^45^. In case of the respiratory syncytial virus (RSV), sequences in the genome rich in G:C nucleotides were recently discovered that promote copy-back RNA synthesis^46^. In this system, these sequences alter polymerase elongation capacity^46^. It is intriguing to speculate that these sequences promote backtracking. While our single-molecule experiments failed to correlate backtracking to specific sequence triggers, consistent with published cell-based recombination studies^14^, analogous to the findings in RSV^46^ the incidence of copy-back RNA synthesis increased in GC-rich regions of template (**Supplementary Fig. 3**).

Our interpretation that the backtracked state and copy-back synthesis contributed to recombination and were of biological relevance was supported by evaluation of RdRp derivatives known to be defective for recombination in cells (Fig. 2)^19^. PV Y275H RdRp is one of the most robust, recombination-defective mutants described^19^, but the molecular basis for the defect is not known for this mutant or the others that have been reported in the PV system or other viral systems^20,43,44^. Here, we show that EV-A71 Y276H RdRp is equally robust and recombination defective in cells (Fig. 2). Both the PV and EV-A71 recombination-defective derivatives retain the ability to sense and respond to errors based on their ability to pause even more frequently (and with longer duration) than WT (Fig. 3B,C and Fig. 4B,C). The increased pausing did not translate to an equal increase in backtracks or copy-back synthesis (Figs. 3E, 4E). The inability to produce the 3’ ssRNA intermediate following backtracking appeared to be caused by an increase in the stability of these enzymes with the 3’-end of nascent RNA-template duplex (Figs. 3G, 4G). We conclude that release of the nascent RNA-template duplex by the RdRp is an obligatory step in converting the paused RdRp elongation complex into a backtracked state with a recombinogenic, single-stranded 3’-end. This level of detail for the mechanism of RdRp-mediated RNA recombination is unprecedented. Several recombination-defective/impaired RdRp variants are known^43,44,47^, and it is possible that the mechanistic basis for the defects will be different. Therefore, evaluation of these derivatives may enable a genetic dissection of the mechanism of template switching and illumination of steps and/or intermediates masked by analyzing WT RdRp.

Given the details on the mechanism of template switching elucidated by this study, it is now clear that the conformational dynamics of the RdRp required to support template switching are substantial. For intramolecular template switching (copy-back synthesis) to occur, the enzyme has to flip to move in the opposite direction and expand the RNA-binding channel to accommodate an additional strand of RNA. For intermolecular template switching (homologous recombination) to occur processively, the enzyme would have to relocate from the site of the backtrack to the new primed-template junction without dissociating into solution. Such polymerase acrobatics is well documented for HIV RT^48–50^. Interestingly, nucleotide analogues also promote substantial changes in the conformational dynamics of HIV RT, as observed here for the EV-A71 RdRp^49^.

T-1106 appears to be unique in its ability to efficiently induce backtracking intermediates especially when compared to structurally similar analogues like ribavirin, a triazole carboxamide^13^. Perhaps, the ability of the T-1106 pseudo base to form a transient basepair within nascent RNA enables priming of copy-back synthesis. The studies reported here are insufficient to provide a definitive explanation, and future biophysical studies of the energetics of T-1106 pseudo base pairing are warranted.

In conclusion, our studies reveal the power of the magnetic tweezers for the characterization of RdRp-catalyzed RNA recombination. The ability to detect an important recombinogenic intermediate and follow the fate of that intermediate provides the first direct mechanistic connection between RNA recombination and copy-back RNA synthesis. We have further elaborated the mechanism of action of the pyrazine carboxamide class of compounds and have discovered that T-1106 functions by promoting formation of the recombinogenic intermediate. This discovery leads us to suggest that this class of compounds may be particularly efficacious against those viruses either known to produce defective viral genomes by copy-back RNA synthesis or associated with high rates of recombination. An important aspect of this class of antiviral compounds is that the barrier to resistance for the virus may be insurmountable. Mutants that were resistant to T-1106-induced template switching remain highly sensitive to the drug, presumably because of the inability to combat the deleterious effects of accumulated mutations (Fig. 6H-J and **Supplementary Fig. 6B**)^40,43,44^. Altogether, our results define inducible intra-and intermolecular template switching (Fig. 6K) as a new viable mechanistic target with broad-spectrum appeal.

## SUPPLEMENTAL INFORMATION

Supplemental Information includes six figures, one table, and the experimental section, and can be found with this article online.

## ACKNOWLEDGEMENTS

Funding to S.-R.S., C.E.C., and N.H.D. was provided by the Human Frontiers Science Program (RPG0011/2015). This research was further funded by National Institutes of Health grant R01AI45818 to C.E.C. A.W. is a recipient of a postdoctoral fellowship from the American Heart Association (18POST33960071). S.R.S. was further supported by grants from the Ministry of Science and Technology (MOST), Taiwan (MOST 107-3017-F-182-001), and the Research Center for Emerging Viral Infections from The Featured Areas Research Center Program within the framework of the Higher Education Sprout Project by the Ministry of Education in Taiwan. We thank Wessel Teunisse and Friso Douma for assistance in single-molecule data collection. We are grateful to Theo van Laar for polymerase purification and synthesis of RNA constructs. We further thank Behrouz Eslami-Mossallam, Martin Depken and David Dulin for fruitful discussions.

## AUTHOR CONTRIBUTIONS

R.J., A.W., K.-M.L., J.J.A., B.S., S.-R.S., C.E.C., and N.H.D. conceived and designed the experiments. R.J. and L.K. performed single-molecule RNA synthesis experiments. R.J. performed the sequence analyses and analyzed the single-molecule data. A.W. and F.F. conducted the cell-based recombination assays, viral genome translation studies, and qPCR of recombinant sequences. I.M. carried out the structural molecular dynamics simulations. K.-M.L. and P.-N.H. executed the *in vivo* mouse infection experiments. A.L.P. and D.A.H. synthesized T-1106. All authors discussed the data. R.J., A.W., C.E.C., and N.H.D. wrote the manuscript.

