## Supplementary Information for "Induced copy-back RNA synthesis as a novel therapeutic mechanism against RNA viruses"

TABLE OF CONTENTS

**Supplementary Figures**

**Figure S1:** Chemical structures of pyrazine-carboxamide base analogues.

**Figure S2:** EV-A71 RdRp exhibits increased pausing upon nucleotides deficiency, but similar forward and reverse RNA synthesis dynamics.

**Figure S3:** EV-A71 copy-back RNA synthesis occurs sequence-independent.

**Figure S4:** T-1106 affects EV-A71 RdRp RNA synthesis dynamics and processivity.

**Figure S5:** T-1106 increases PV RdRp pausing and backtracking.

**Figure S6:** T-1106 dose response of EV-A71 WT and Y276H variant full-length viruses.

**Supplementary Tables**

**Table S1:** Single molecule experimental parameters and measurement statistics.

**Materials and Methods**

### SUPPLEMENTARY FIGURES

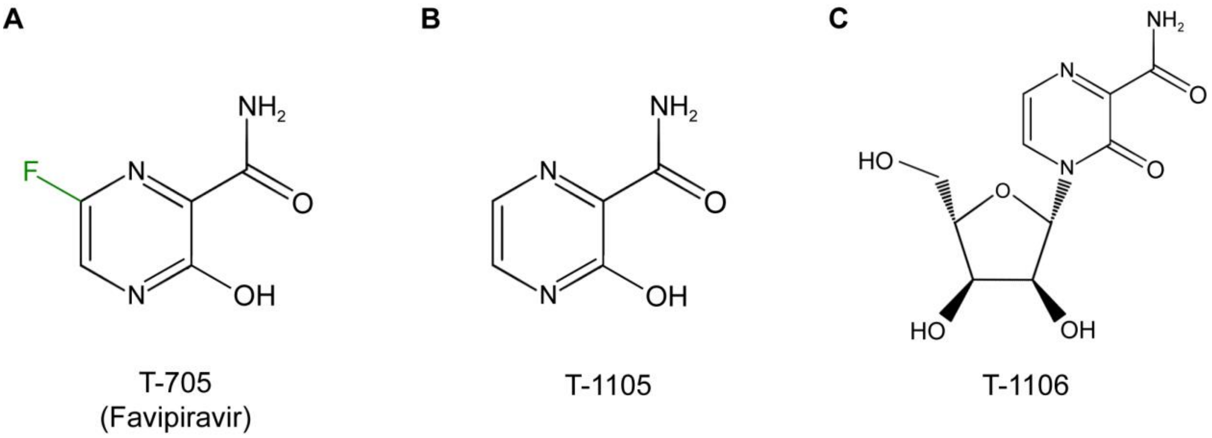

**Supplementary Figure 1. Chemical structures of pyrazine-carboxamide base analogues.** (**A**) Nucleobase T-705, also known as favipiravir. (**B**) The defluorinated T-705 nucleobase-analogue, denoted T-1105. (**C**) The nucleoside version of T-1105, denoted T-1106.

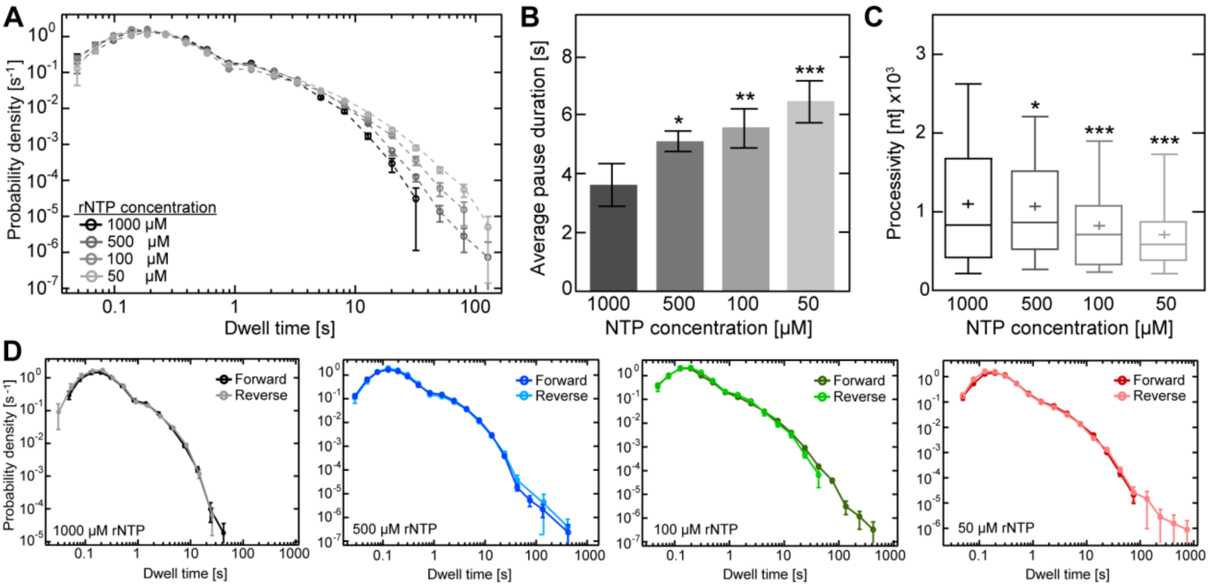

**Supplementary Figure 2. EV-A71 RdRp exhibits increased pausing upon nucleotides deficiency, but similar forward and reverse RNA synthesis dynamics.** (**A**) Dwell time distributions of EV-A71 wild type RdRp RNA synthesis dynamics at different nucleotide concentrations (50 µM - 1 mM). Dwell time window was set to 4 nt and the error bars (±SD) result from bootstrapping with 1,000 iterations. (**B**, **C**) Comparison of extracted quantitative pause values during RNA synthesis upon nucleotide deficiency shown in **Fig. 1**, such as (**B**) average (±SEM) pause duration, and (**C**) RNA synthesis processivity. (**D**) Superimposed dwell time distributions of forward (dark colors) and reverse RNA chain elongation (light colors) at different rNTP concentrations (50 µM - 1 mM). Statistical analyses were performed using one-way analysis of variance (ANOVA) with comparative Tukey post-hoc test (significance levels $\alpha$: *** = 0.001; ** = 0.01; * = 0.05).

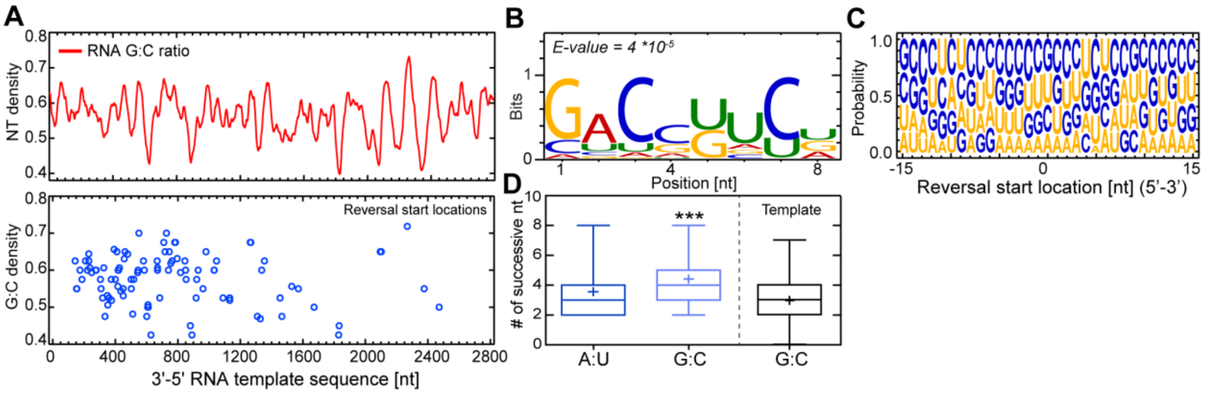

**Supplementary Figure 3. EV-A71 copy-back RNA synthesis occurs sequence-independent**. (**A**) Analysis of sequences of detected template-switching events (lower panel, blue circles) did not exhibit any hotspots in respect to the RNA template sequence (upper panel, red line) where copy-back RNA synthesis predominantly occurs. The G:C density at observed copy-back RNA synthesis locations (lower panel, blue circles) did not reflect any apparent correlation. (**B**) With the absence of apparent copy-back RNA synthesis hotspots, the obtained sequences were subject to *de novo* sequence motif search using MEME, a multiple sequence alignment algorithm that searches for conserved sequence motifs[69]. The computed sequence conservation at locations of reversal events show a bit score <1 and a probability (E-value) strongly below the 95% confidence interval threshold. Both results reveal no evidence of a predominant sequence motif as trigger. (**C**) Sequence logo of position-dependent nucleotide abundance within the copy-back RNA synthesis sequences showed that guanosines and cytosines are most abundant, which can hamper effective melting of the RNA duplex during RNA synthesis, resulting in an increase of RdRp to pause and arrest. In agreement, (**D**) a higher abundance of successive G:C (light blue) nucleotides within the copy-back RNA synthesis sequences is found compared to both the average amount encountered in the ssRNA template (black) and successive A:U (light blue). Statistical analysis were performed using one-way analysis of variance (ANOVA) with comparative Tukey post-hoc test (significance level $\alpha$: *** = 0.001).

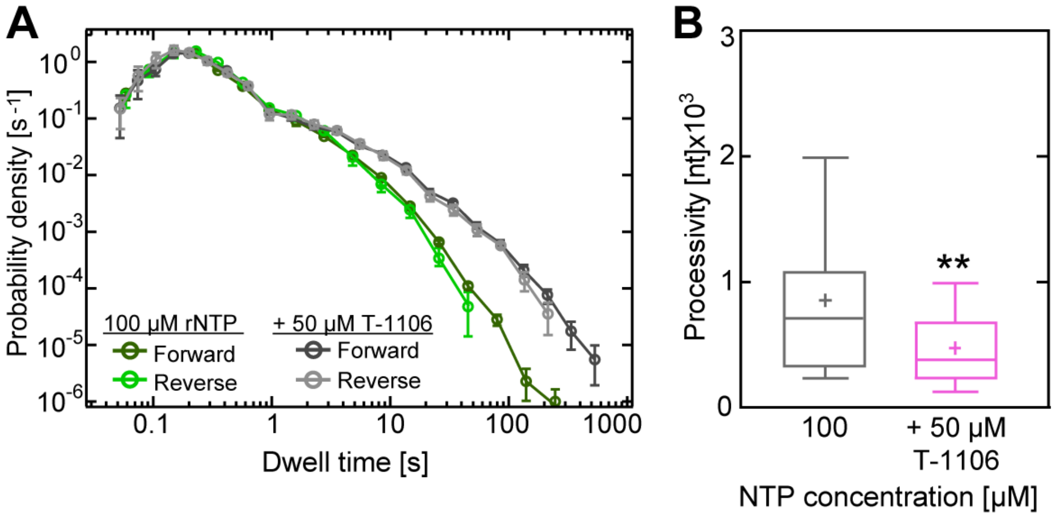

**Supplementary Figure 4. T-1106 affects EV-A71 RdRp RNA synthesis dynamics and processivity.** (**A**) Superimposed dwell time distributions of forward (dark color) and reverse (light color) RdRp translocation in presence (grey) and absence (green) of T-1106. (**B**) T-1106 (magenta) reduces significantly EV-A71 RNA synthesis processivity. Statistical analysis consisted of unpaired, two-tailed t-tests (significance level P: ** $\leq$ 0.01).

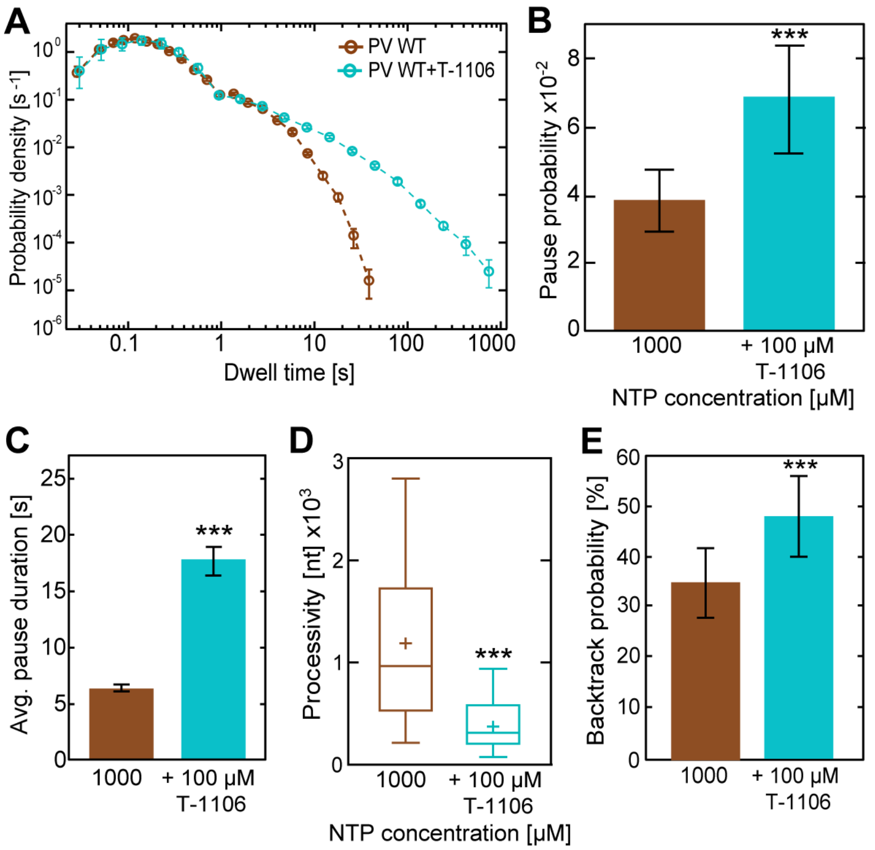

**Supplementary Figure 5. T-1106 increases PV RdRp pausing and backtracking.** (**A**) Superimposed dwell time distributions of PV WT RdRp (brown) exhibit increased pausing probability and duration in presence of T-1160 (cyan). Dwell times are associated with polymerases advancing 4 nt. The error bars represent the estimate of the standard deviations via bootstrapping with 1,000 iterations. (**B-E**) In presence of T-1106, wild type poliovirus RdRp exhibits significantly higher (**B**) average (±SD) pausing probability of (**C**) extended apparent duration (AVG ±SEM) during RNA synthesis, leading to a decreased (**D**) processivity but higher (**E**) backtracking probability. Statistical analysis consisted of unpaired, two-tailed t-tests (significance level P: *** ≤ 0.001; * ≤ 0.05).

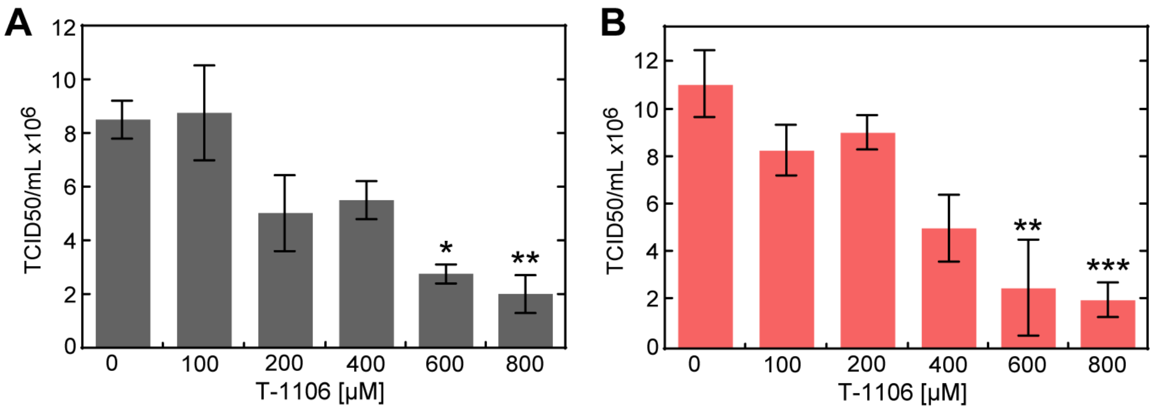

**Supplementary Figure 6. T-1106 dose response of EV-A71 WT and Y276H variant full-length viruses.** RD cells infected with (**A**) EV-A71 wild type and (**B**) Y276H donor genomes at MOI 1 in the presence of increasing concentration of T-1106. Following total cytopathic effect (CPE), cell culture supernatants were used to quantify virus by $TCID_{50}$. Statistical analyses were performed using one-way analyses of variance (ANOVA) with comparative Tukey post-hoc test (significance levels α: *** = 0.001; ** = 0.01; * = 0.05).

### SUPPLEMENTARY TABLES

**Supplementary Table 1: Single molecule experimental parameters and measurement statistics.** Table of experimental parameters, such as NTP and T-1106 concentrations, as well as dwell times described as statistical measure for all experiments performed in this study. The number of nucleotides is an alternative statistical measure that reflects the total length of synthesized RNA monitored from pooled trajectories. The applied force was hold constant at 25 pN for all experiments.

| **RdRp** | **EV-A71 C2-MP4** | | | | |
|---|---|---|---|---|---|
| **[NTP]** | 1 mM | 500 µM | 100 µM | 50 µM | 100 µM |
| **[NA]** | - | - | - | - | 50 µM T-1106 |
| **Dwell times** | **9981** | **16356** | **13430** | **10042** | **6570** |
| **Nucleotides** | 39.9 kb | 65.4 kb | 53.7 kb | 40.1 kb | 26.3 kb |

| **RdRp** | **EV-A71 Y276H** | **PV WT** | **PV WT** | **PV Y275H** |
|---|---|---|---|---|
| **[NTP]** | 1 mM | 1 mM | 1 mM | 1 mM |
| **[NA]** | - | - | 100 µM T-1106 | - |
| **Dwell times** | **4881** | **5715** | **4839** | **3836** |
| **Nucleotides** | 19.5 kb | 22.9 kb | 19.4 kb | 15.3 kb |

### MATERIALS AND METHODS

### MATERIALS

#### KEY RESOURCES LISTING

| REAGENT or RESOURCE | SOURCE | IDENTIFIER |
|---|---|---|
| **Antibodies** | | |
| Digoxigenin antibodies | Roche | Cat# 11333089001 |
| **Bacterial and Virus Strains** | | |
| SURE Competent Cells | Agilent technologies | Cat#200238 |
| **Chemicals, Peptides, and Recombinant Proteins** | | |
| ApU dinucleotide | IBA Lifesciences GmbH | Cat#0-31004 |
| Biotin-16-dUTP | Roche | Cat#11093711103 |
| Digoxigenin-11-dUTP | Roche | Cat#11093681103 |
| rNTPs | GE Healthcare | Cat#27-2025-01 |
| Streptavidin-coated superparamagnetic beads | Thermo Fischer | Cat#65001 |
| T-1106 triphosphate | Blake Petersen Lab | N/A |
| Ribavirin triphosphate | Jena Bioscience | Cat#NU-1105L |
| Superase RNase inhibitor | ThermoFischer | Cat#AM2694 |
| **Critical Commercial Assays** | | |
| Ribomax large scale RNA production kit | Promega | Cat#P1300 |
| RNeasy MinElute cleanup kit | Quiagen | Cat#74204 |
| **RNA Oligonucleotides** | | |
| 5’-AACUGUUGGUGUACGCGAAAGCGU | GE Healthcare Dharmacon, Inc. | |
| **DNA Oligonucleotides** | | |
| 5’-taatacgactcactataggatcgccaagattagcggatcctacctgac | Biolegio | AB-For |
| 5’-ggttaacctcaacttccatttcc | Biolegio | AB-Rev |
| 5’-cccctcgaggggaaaaaaaaaaaaaccgtatgacgctggaag | Biolegio | CD-For |
| 5’-taatacgactcactataggccggacgtttcggatcttccgacatgcgc | Biolegio | CD-Rev |
| 5’-aagattagcggatcctacctgac | Biolegio | Bio-For |
| 5’-bio-taatacgactcactataggaacggcttgatatccactttacg | Biolegio | Bio-Rev |
| 5’-agcgtaaaattcagttcttcgtggcg | Biolegio | Dig-For |
| 5’-dig-aatacgactcactatagggctaccggttaacctcaacttccatttcc | Biolegio | Dig-Rev |
| 5’-tgccattcagggactgccgatgtcggtgcagccg | Biolegio | SP-For |
| 5’-taatacgactcactataggagcgccgcttccatgtcctggaacgct | Biolegio | SP-Rev |
| 5’-acgttctcagtgcggactgtag | IDT | MP4-3156-Fwd |
| 5’-ccttggaaaagagcttc | IDT | C2/4-5057-Rev |
| 5’-gaagctcttttccaagg | IDT | C2/4-5057-Fwd |
| 5’-ctggttataacaaatttaccc | IDT | C2/4-7384-Rev |
| 5’-atcaatcacacccatcatgtgcatcgcaataaaacttattg | IDT | Y276H-Fwd |
| 5’-caataagttttattgcgatgcacatgatgggtgtgattgat | IDT | Y276H-Rev |
| 5’-acgtagcccagcgcgtcggccg | IDT | Eag-I-Rev |
| 5’-agcagtgctgactagtaaagacc | IDT | SpeI-Fwd |
| 5’-cggcagcccagaagaact | IDT | qPCR-Fwd |
| 5’-gccaccctatctccctgga | IDT | qPCR-Rev |
| 5’-FAM-tcaccatgaagttgtgtaaggatgcta-BHQ | Biosearch Technologies | qPCR-Probe |
| **Recombinant DNA** | | |

| | | |
|---|---|---|
| Plasmid pBB10 | Petrushenko *et al.*, 2006[52] | pBB10 |
| pSumoEV-A71-3D | This study | |
| pSumoEV-A71-3D-Y276H | This study | |
| pSumoPV-3D | Arnold J.J. & Cameron C.E., 2000[23] | |
| pSumoPV-3D-Y275H | Acevedo *et al*., 2018[19] | |
| pEV-A71-MP4 | Wang *et al*., 2004[51] | |
| pEV-A71-Y276H | This study | |
| pEV-A71-C2-replicon | Woodman *et al*., 2018[14] | |
| pEV-A71-MP4-Δ3D | Woodman *et al.*, 2018[14] | |
| **Experimental Models: Cell Lines** | | |
| RD cells | ATCC | Cat#CCL-36 |
| **Software and Algorithms** | | |
| MatLab R13 | MathWorks Inc. | www.mathworks.com |
| Igor Pro 6.37 | Wavemetrics | www.wavemetrics.com |
| LabView 2011 | National Instruments | www.ni.com |
| ImageQuant | GE Healthcare Life Sciences | www.gelifesciences.com |
| ImageLab | BioRad | www.bio-rad.com/en-ch/product/image-lab-software |

### CONTACT FOR REAGENT AND RESOURCE SHARING

### METHODS

#### Cell culture

Adherent monolayers of human embryonic rhabdomyosarcoma (RD) cells were grown in Dulbecco’s Modified Eagle Medium (DMEM). Media was supplemented with 100 U/ml penicillin, 100 µg/ml streptomycin, and 10% Heat Inactivated (HI)-FBS. All cells were passaged in the presence of trypsin-EDTA. Cell were maintained at 37°C/5% $CO_2$.

#### Plasmids

The mouse adapted EV-A71 C2-MP4 infectious clone was kindly provided by Dr. Jen-Reng Wang (Cheng Kung University, Taiwan) and modified by insertion of a ribozyme sequence between the T7 promoter and viral genome sequence in a pBR-derived plasmid[52,53].The EV-A71 C2 replicon was modified from a previously described EV-A71 C2-2231 replicon by addition of a T7-ribozyme and polyA sequence inserted at the 5’ and 3’ end of the replicon sequence in a pBR-derived plasmid[54]. The EV71Δ3D template was constructed from the full-length EV-A71 C2-MP4 infectious clone by removal ~800 nt between the blunt cutting restriction sites (*Sca*I and *Nru*I) within the 3D$^{pol}$ coding region. The Y276H mutant replicon and infectious clone were constructed by using site-directed mutagenesis. The sequences of the primers (Integrated DNA Technologies, Inc.; IDT) used for the plasmid construction are shown in the Key Resources Table.

**Purification, 5′-$^{32}$P Labeling, and Annealing of sym/sub**

RNA oligonucleotides were purified, labeled, and annealed as described previously Arnold JJ & Cameron CE[23]. Poliovirus RNA-dependent RNA polymerase (3D$^{pol}$): Assembly of stable, elongation-competent complexes by using a symmetrical primer-template substrate (sym/sub).

**Expression and purification of Enterovirus A-71 RdRp**

Mutation of the Y276 codon was performed by standard PCR mutagenesis. Expression and purification of WT and mutant 3D$^{pol}$ enzymes followed previous procedures with some minor modifications[31,55,56]. 3D$^{pol}$ is expressed as a fusion protein to SUMO and an N-terminal polyhistidine tag that increases protein production, eases purification and allows for production of 3D$^{pol}$ with the naturally occurring Gly1[55,56].

Protein purification: buffer B (100 mM potassium phosphate, 500 mM NaCl, 5 mM imidazole, 5 mM β-mercaptoethanol, 60 μM $ZnCl_2$, 20% w/v glycerol, pH 8.0) and buffer C (100 mM potassium phosphate, 500 mM NaCl, 60 μM $ZnCl_2$, 5 mM β-mercaptoethanol, 20% w/v glycerol, pH 8.0) were prepared. Cell pellets were resuspended in 50 ml lysis buffer (50 ml buffer B, 1.4 μg/ml pepstatin A, 1 μg/ml leupeptin, 1 mM PMSF, 0.1% N-P40) and subjected to sonication. Cell lysates were centrifuged at 30,000 g and 4°C for 30 min. Supernatant was applied to Ni-NTA (Invitrogen) columns pre-equilibrated with buffer C1 (buffer C, 5 mM imidazole and 0.1% N-P40). The resin was washed with three bed volumes each of buffer C1 and buffer C2 (buffer C, 5 mM imidazole), and protein was eluted using high imidazole buffers C3 (buffer C, 50 mM imidazole) and C4 (buffer C, 500 mM imidazole). The polyhistidine tag and SUMO protein domain were cleaved from 3D$^{pol}$ using the protease Ulp1. Protein solutions were dialyzed against 80 mM Tris-HCl, 500 mM NaCl, 20% w/v glycerol, 10 mM β-mercaptoethanol, 60 μM $ZnCl_2$, pH 8.0 overnight (optimal buffer for protease cleavage), and then dialyzed against 100 mM potassium phosphate, 20% w/v glycerol, 10 mM β-mercaptoethanol, 60 μM $ZnCl_2$, pH 8.0 for 2–3 hours. A second Ni-NTA column was used to separate the purified 3D$^{pol}$ from the cleaved polyhistidine tag/SUMO domain, using procedures identical to the first Ni-NTA column. For WT and Tyr276His 3D$^{pol}$, additional phospocellulose and Q-sepahrose columns were used to ensure protein solutions were free of trace contaminants of nuclease and phosphatase activities[55,56]. Proteins were >95% homogeneous as estimated by coomassie-blue staining of SDS PAGE gels. Protein is stable at 4°C for 3-4 months under high salt conditions (80 mM Tris-HCl, 500 mM NaCl, 20% w/v glycerol, 10 mM β-mercaptoethanol, 60 μM $ZnCl_2$, pH 8.0).

***In vitro* RNA synthesis, cell transfection, and recombinant virus quantification**

The EV-A71 C2 replicon and C2-ΔIRES-replicon were linearized with *Sal*I. The EV-A71-MP4, EV71Δ3D cDNA were linearized with *Eag*I. All linearized cDNA was transcribed *in vitro* using T7 RNA Polymerase treated with 2U DNAse Turbo (ThermoFisher) to remove residual DNA template. The RNA transcripts were purified using RNeasy Mini Kit (Qiagen) before spectrophotometric quantification. Purified RNA (amounts as specified elsewhere) in RNase-free $H_2O$ were transfected into cell lines using TransMessenger (Qiagen). The mixture was incubated according to the manufacturer's instructions and added to RD cell monolayers in 12-well tissue culture plates. Virus amount was quantified by plaque assay. Briefly, media supernatant and cells were harvested at time-points post transfection (specified in the main text), subjected to three freeze-thaw cycles and clarified. Supernatant was then used on fresh RD cells in 12-well plates, virus infection was allowed to continue for 30 min. Media was then removed, and cells were subjected to 2x PBS (pH 7.4) washes before a 1% (w/v) agarose-media overlay was added. Cells were incubated for 3-4 days and then fixed and stained with crystal violet for virus quantification.

**Single-step growth curve for EV-A71 WT and Y276H full-length virus**

RD cells in 12-well plates were infected by each virus at a MOI of 0.1 in triplicate in serum-free media. One hour later, cells were extensively washed by PBS and refreshed in 10% serum-containing media. Virus was harvested at different time-points post infection and the virus yield was quantified by plaque assay.

**Real time qPCR analysis**

Viral RNA was isolated with QiaAmp viral RNA purification kit (Qiagen), as recommended by the manufacturer. The real time qPCR analysis was performed by the Genomics Core Facility of The Pennsylvania State University. DNase-treated RNA was reverse-transcribed using the High Capacity cDNA reverse transcription kit (Applied Biosystems, Foster City, CA, USA) and the protocol provided with the kit. Quantification by real time qPCR was done by adding 10 or 20 ng of cDNA in a reaction with 2x TaqMan Universal PCR Master Mix (Applied Biosystems, Foster City CA) in a volume of 20 µl, with primers 5′-CGGCAGCCCAGAAG AACT-3′ (forward) and 5′-GCCACCCTATCTCCCTGGAT-3′ (reverse) and probe 5′-[6-Fam]-TCACCATGAAGTTGTGTAAGGATGCTA-3′ in a 7300 real time qPCR machine (Foster City, CA, USA). A standard curve was generated using *in vitro* transcribed RNA.

**Luciferase assays**

Supernatant was removed from transfected cell monolayers, and cells were briefly washed with PBS and lysed using 100 µl 1x Glo Lysis Buffer (Promega®) per well in a 12-well plate. The oxidation reaction was catalyzed by the addition of 10 µl cell lysate to 10 µl room temperature *Bright-Glo* Luciferase Assay System (Promega®) substrate. Luciferase activity was measured using a luminometer with values normalized to protein content of the extract using a protocol as previously described[57].

**Generation and infection of transgenic human hSCARB2–expressing mice**

Transgenic mice expressing hSCARB2 were generated as described previously[58], and kindly provided by Dr. Satoshi Koiki (Tokyo Metropolitan Institute of Medical Science, Japan). For infection, $2x10^7$ genome copies of wild type and Y276H mutant EV-A71 C2-MP4 viruses were intragastrically inoculated into 21-day old mice using metallic gastric tubes. The inoculated mice were monitored daily for clinical disease symptoms, reflected by the disease scores defined as: 1. jerky movement; 2. paralysis of one hind leg; 3. paralysis of both hind legs; 4. death.

**Bulk RdRp turnover experiments**

1 µM WT and YH variants of poliovirus or EV-A71 C2-MP4 RdRp were incubated with $32^{P}$-labelled 20 µM RNA primer-template duplex (sym/subU) and 500 µM rNTPs. At various time points (described in text) the reaction was quenched by the addition of 500 mM EDTA and 35% formamide. Products were analyzed *via* denaturing polyacrylamide gel electrophoresis.

**Denaturing polyacrylamide gel electrophoresis**

Quenched reactions were mixed with an equal volume of loading buffer (75% formamide, 0.025% bromophenol blue and 0.25% xylene cyanol) and heated to 70°C prior to loading on a denaturing PAGE gel. Products were resolved from substrates by denaturing polyacrylamide gel electrophoresis (18.5% acrylamide, 1.5% bisacrylamide, 1x TBE buffer, 7 M Urea). Electrophoresis was performed in 1x TBE at 90 W. Gels were visualized using a PhosphorImager and quantified using ImageQuant software (GE Healthcare).

**RNA constructs for single-molecule RNA synthesis experiments**

The RNA template used in our single-molecule assay is analogous to the sequences used in previous studies of PV and **Φ**6 RdRp RNA synthesis kinetics[13,16]. The dNA tether construct consists predominantly of a dsRNA, assembled by hybridization of a 2.8 kb template ssRNA to a 4.1 kb complementary strand, and two ~ 500 bases ssRNA strands containing either biotin or digoxigenin for the tethering between magnetic beads and the surface, as previously described in detail[13]. In contrast to the previously used hairpin for Poliovirus RdRp RNA synthesis initiation, terminating the 3' end of the template, we used for the hairpin structure 24 bases with the following sequence: 5'-AACUGUUGGUGUACGCGAAAGCGU-3'. Essentially, the hairpin mimics a primer and promotes primer-dependent RNA synthesis initiation[21].

The RNA constructs were assembled by first subjecting Plasmid pBB10 to PCR amplification using primers AB-For, AB-Rev, CD-For, CD-Rev, Bio-For, Bio-Rev, Dig-For, Dig-Rev, SP-For, and SP-Rev, listed in the Key Resource Listing. Single-stranded RNA sequences were produced from these amplicons *via in vitro* run-off transcription using T7 RNA polymerase from the Ribomax large-scale RNA production system (Promega). Transcription reactions for the AB, CD and SP ssRNA fragments contained 500 ng DNA amplicon, 10 µl T7 buffer, 5 µl T7 polymerase, 1 µl 100 mM CTP, and 1 µl of 100 mM for all other NTPs (ATP, UTP, GTP) in a reaction volume of 50 µl. For the synthesis of BIO ssRNA, the UTP amount was reduced to half and the reaction mix was supplemented with 4.7 µl of 10 mM biotin-16-UTP (Roche), while for DIG ssRNA the UPT amount was reduced only to 0.63 µl with supplement of 3.7 µl or 10 mM digoxigenin-11-UTO (Roche). The synthesized ssRNA fragments were purified using an RNeasy MinElute cleanup kit (Qiagen) and eluted in 1 mM sodium citrate buffer (pH 6.4).

These different ssRNA fragments were then assembled into a dsRNA construct for the single-molecule studies conducted on Poliovirus and Enterovirus A-71 RdRp *via* hybridization. The ssRNA fragments were first mixed in equimolar ratio in 200 µl 0.5x SSC buffer, with the exception of the biotin (BIO) and digoxigenin (DIG)-enriched handles, which were added in four times molar excess in respect of the AB strand (1 µg AB, 1.4µg CD, 360 ng SP, 450 ng BIO, and 516 ng DIG). Afterwards, the RNA mix was heated for 1 h to 65°C, cooled with a rate of 0.24°C/min down to 25°C. The final dsRNA construct was purified with RNeasy MinElute cleanup kit and eluted in 1 mM sodium citrate. All RNA concentrations were photometrically determined (Nanodrop).

**Magnetic tweezers experimental configuration**

The magnetic tweezers implementation used in this study has been described previously[13]. Briefly, light transmitted through the sample was collected by a 50x oil-immersion objective (CFI Plan 50XH, Achromat, 50x, NA = 0.9, Nikon) and projected onto a 12 megapixel CMOS camera (#FA-80-12M1H, Falcon2, Teledyne Dalsa) with a sampling frequency of 50 Hz. The applied magnetic field was generated by a pair of vertically aligned permanent neodymium-iron-boron magnets (Webcraft) separated by a distances of 1 mm, suspended on a motorized stage (#M-126.PD2, Physik Instrumente) above the flow cell. Image processing of the collected light allows tracking the real-time position of both surface attached reference beads and superparamagnetic beads coupled to the dsRNA constructs in three dimensions over time. The bead *x*, *y*, *z* position tracking was achieved using a cross-correlation algorithm realized with custom-written software in LabView (2011, National Instruments Corporation)[59]. Bead positions were determined with spectral corrections to correct for camera blur and aliasing[59].

**Single-molecule RdRp RNA synthesis assay**

The flow cell preparation used in this study has been described in detail elsewhere[13]. In brief, polystyrene reference beads (#17133, Polysciences GmbH) of 1.5 µm in diameter were diluted 1:1500 in PBS buffer (pH 7.4; Sigma Aldrich) and then adhered to the nitrocellulose-coated (Invitrogen) surface of the flow cell. Afterwards, digoxigenin antibodies (Roche Diagnostics) at a concentration of 0.1 mg/ml were incubated for 1 hour within the flow cell, following a 2-hour incubation of 10 mg/ml BSA (New England Biolabs) diluted in PBS (pH 7.4) buffer. After washing with PBS buffer, 100 µl of streptavidin-coated superparamagnetic beads (DynaBeads, LifeTechnologies; prior diluted 1:400 from stock) with a diameter of 1.5 µm were added resulting in the attachment of the beads to the surface-tethered dsRNA constructs. Afterwards, unbound beads were washed out with PBS buffer.

The preparation of RdRp:RNA ternary complexes was also performed as described previously[13,16]. Briefly, 1 µM RdRp in 100 µl EV buffer (50 mM HEPES, 5 mM MgCl2, 125 µg/ml BSA, 1 mM DTT, 1 U Superase RNase inhibitor, pH 6.6), supplemented with 600 µM ATP, 600 µM CTP, and 1.2 mM ApC, was flushed into the flow cells containing the dsRNA constructs. Stalled RdRp complexes were formed during 20 minutes of incubation at room temperature. Afterwards, the flow cell was washed with EV buffer and the RNA chain elongation was re-initiated by adding all four rNTP in an equimolar concentration of 1 mM, unless stated otherwise, and in presence or absence of nucleotide analogues. The single-molecule measurements were conducted for 2 hours at constant pulling forces of 25 pN at 24°C with a camera acquisition rate of 50 Hz.

**Molecular Dynamics simulations (MD)**
The starting coordinates for the all-atom MD simulations systems prepared by the accessory program "tleap" of AMBER18 suite[70]. The crystal structure 3N6L of EV71 RdRp was used to prepare the starting coordinates for MD simulations of the WT system[60]. The Y276H mutant system prepared by *in silico* replacement of Tyr at position-276 by a His; the steric clashes generated in the mutant system removed by subsequent energy minimization and equilibration. The WT PV RdRp was investigated previously by all-atom MD simulations (150 ns), using the 1RA6 structure as starting coordinates for the MD simulations[29,31].

For the performed MD simulations, the protein parameters of Amber14SB force field was used during calculations[62]. All MD simulations were performed in explicit water (TIP3P model), imposing a minimal distance of 12 Å between the edge of the solvent box and any protein atom[63]. Calculations of the non-bonded interactions used a cutoff radius of 9 Å with periodic boundary conditions applied; particle mesh Ewald method was used to treat electrostatic interactions[64]. The SHAKE algorithm was employed to constrain hydrogens bonded to heavy atoms[65]. The simulations were performed by first relaxing the systems in two cycles of energy minimization using SANDER program. Subsequently, the systems were slowly heated to 300 K using the parallel version PMEMD under NVT conditions (constant volume and temperature); Langevin dynamics with collision frequency ($\gamma = 2$) was used to regulate temperatures[66]. The heated systems were then subjected to equilibration by running 100 ps of MD simulations under NPT conditions (constant pressure and temperature) with 1 fs integration time steps. MD trajectories were collected over 200 ns at 1 ps interval and 2 fs integration time steps. Analyses of the trajectories from MD simulations were performed using CPPTRAJ program[67]. MD simulations were carried out on a multi-GPU workstation with 32-core processor (AMD Ryzen Threadripper 2990WX) and four Nvidia GTX 1080 Ti graphics cards.

### QUANTIFICATION AND STATISTICAL ANALYSIS

**Single-molecule data processing**
RNA synthesis trajectories were processed using custom-written Igor v6.37 and MatLab R2013b-based custom-written scripts. The measured z-position of RdRps during the transcription process were converted to transcribed RNA product as a function of time, using the empirically determined force-extension relationships for dsRNA and ssRNA molecules under the employed buffer conditions[13,16]. To reduce the effect of Brownian noise in the applied statistical analyses, all measured elongation trajectories were filtered to 1 Hz using a sliding mean average filter.

**Statistical dwell time analysis of single RdR elongation trajectories**
The stochastic RNA synthesis dynamics of EV-A71 and PV RdRp were quantitatively assessed by a statistical analysis of elongation and pausing using a recently described bias-free dwell-time analysis[13,16,68]. Using this approach, the times needed for RdRp to elongate through consecutive dwell time windows of four nucleotides - defined as *dwell times* - were determined for all

measured RdRp trajectories under the same conditions to construct dwell time probability distributions (e.g. **Supplementary Fig. 2A**). The dwell times were bootstrapped 1,000 times to estimate the standard deviation and confidence intervals of the distributions[13,68]. All dwell-time distributions qualitatively exhibit the same features, such as a peak at short time scales (~ 200 ms) and a tail of gradually decreasing probability in time scales ranging between 1 and 1,000 s. While the peak reflects fast kinetic processes consisting of nucleotide addition, NTP hydrolysis, $PP_i$ release, and translocation, the tail originates from off-pathway pauses, consistent with the previous observations for the PV RdRp[13].

The quantitative description of the duration and probability of pauses during RNA synthesis derive from these dwell time distributions. To calculate the average pause probabilities (±SD), the dwell time distributions were integrated starting from a chosen threshold of 3 s, where single pauses are detectable within a dwell time window of 4 nt, similar to the value obtained in our previous single-molecule study of PV RdRp dynamics[13]. The apparent lifetime of these pauses is determined by averaging all dwell times (±SEM) exceeding this threshold of 3 s.

The probabilities of EV-A71 copy-back RNA synthesis (also referred as *reversals*) and PV backtracking were determined by dividing their occurrence by the total amount of RNA synthesis trajectories per pooled data set.

The enzyme processivites reflect the measured length of synthesized RNA chains (in nucleotides) for each measured RdRp defining their termination event, and the average velocity (± SD) resulted from dividing individual RdRp processivities by their active RNA synthesis duration.

**Sequence analysis single-molecule reversal event locations**

The sequence analysis was performed similar to the analysis used in the previously published study of EV-A71 recombinants yielding from transfected cell-based assays[14]. Here, the locations at which template switching was observed were extracted from single-molecule experiments and subject to sequence analysis. We used a sequence window of 30 nt (-15 to +15 nt around the detected locations) for the ssRNA template. This window size was determined based on two parameters. First, in a recently reported EV-A71 cell-based recombination assay, ‘copy-choice’ recombination occurred in regions with a sequence homology of between 5-11 nt between the two parental template strands[14]. Second, the accuracy of the used assay is limited by the stiffness of the ssRNA construct and the corresponding degree of Brownian noise which accounts for a resolution of ±10 nt. The window size of 30 nt therefore guarantees to comprise any possible sequence motif that could be encountered *in vivo*. From these extracted location sequences, we calculated G:C and A:U densities, as well as the amount of successive G|C or A|U nucleotides. *De novo* sequence motif search for these location sequences were performed using the MEME suite (Version 5.0.1)[69] with 1$^{st}$-order background modelling and variable sequence motif width between 4-10 nt.

### DATA AND SOFTWARE AVAILABILITY
